# Historical trade routes for diversification of domesticated chickpea inferred from landrace genomics

**DOI:** 10.1101/2021.01.27.428389

**Authors:** Anna A. Igolkina, Nina V. Noujdina, Maria G. Samsonova, Eric von Wettberg, Travis Longcore, Sergey Nuzhdin

## Abstract

According to archaeological records, chickpea (*Cicer arietinum*) was first domesticated in the Fertile Crescent 10 thousand years ago. Its subsequent diversification in South Asia, Ethiopia, and the Western Mediterranean, however, remains obscure and cannot be resolved using only archeological and historical evidence. In particular, chickpea has two market types: ‘desi’, which has a similar flower and seed coat color to chickpea’s wild relatives; and ‘kabuli’, which has light-colored seed, and is linguistically tied to Central Asia but has an unknown geographic origin.

Based on the genetic data from 421 chickpea landraces from six geographic regions, we tested complex historical hypotheses of chickpea migration and admixture on two levels: within and between major regions of cultivation. For the former, we developed popdisp, a Bayesian model of population dispersal from a regional center towards sample locations, and confirmed that chickpea spread within each region along trade routes rather than by simple diffusion.

For the latter, migration between regions, we developed another model, migadmi, that evaluates multiple and nested admixture events. Applying this model to desi populations, we found both Indian and Middle Eastern traces in Ethiopian chickpea, suggesting presence of a seaway from South Asia to Ethiopia — and the cultural legacy of the Queen of Sheba. As for the origin of kabuli chickpeas, we found significant evidence for an origin from Turkey rather than Central Asia.

## Introduction

The genetic variation of species reflects evolutionary history. The history of a domesticated species is inextricably linked with human history and we can learn much about one from studying the other. Reconstructing the spread of cultigens reveals the history of both plant and human and has the potential to improve modern genomics-assisted breeding schemes.

Chickpea (*Cicer arietinum* L.) is an important source of high-quality protein (Abbo et al., 2003a), ranked third among legumes in terms of grain production (Jain et al., 2013). It is extensively cultivated in India, West Asia, Eastern Africa, and the Mediterranean Basin, but how it reached these regions, and its subsequent admixture history is not well-understood. Limiting factors in reconstructing chickpea domestication history include: (1) lack of whole-genome sequences from ancient chickpea, (2) reduced genetic diversity in cultivars due to domestication bottlenecks, (3) the replacement of locally evolving landraces with modern commercial varieties (Abbo et al., 2003a). The most suitable material for studying chickpea domestication is the historical germplasm collection made by Vavilov in the 1920s-1930s, stored at the N.I. Vavilov All Russian Institute of Plant Genetic Resources (VIR). This collection currently contains 3380 chickpea accessions, almost half of which represent pre-Green Revolution landraces with known geographical origin (Figure 1a). Vavilov not only established this unique collection, but also identified several “centers of origin” (or diversity) of crop plants (Vavilov, 1926) (Figure 2a). For chickpea, centers of diversity include six regions (van der Maesen, 1984; Vavilov, 1951), which we will denote by the nearest contemporary country: Turkey, Uzbekistan, India, Lebanon, Morocco, and Ethiopia. We assembled a panel of 421 chickpea landraces which represent these regions (Figure 1a) and tested historical hypotheses of chickpea diversification based on genotyping at 2579 loci.

**Figure 1.**
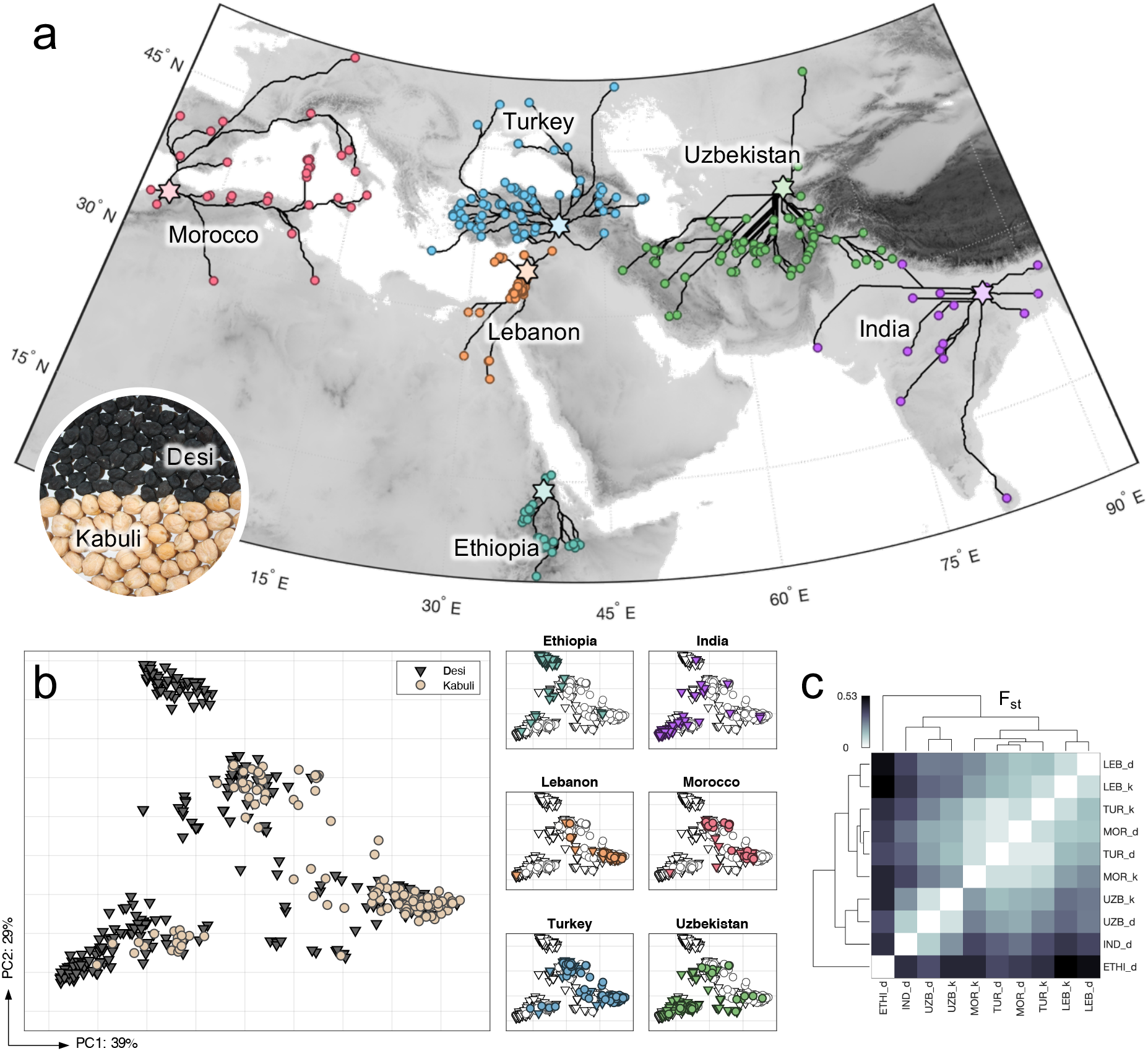
(a) Sampling locations of chickpea accessions (circles) and estimated trade routes from the centers of clusters (stars) to locations. Each net of routes represents a binary tree. Photo shows the morphological differences between seeds of desi and kabuli chickpea types. (b) PCA plots for accession based on SNP data separately colored by chickpea type (left) and by regions (right). (c) Mean pairwise Fst comparison of 10 chickpea subpopulations.

**Figure 2.**
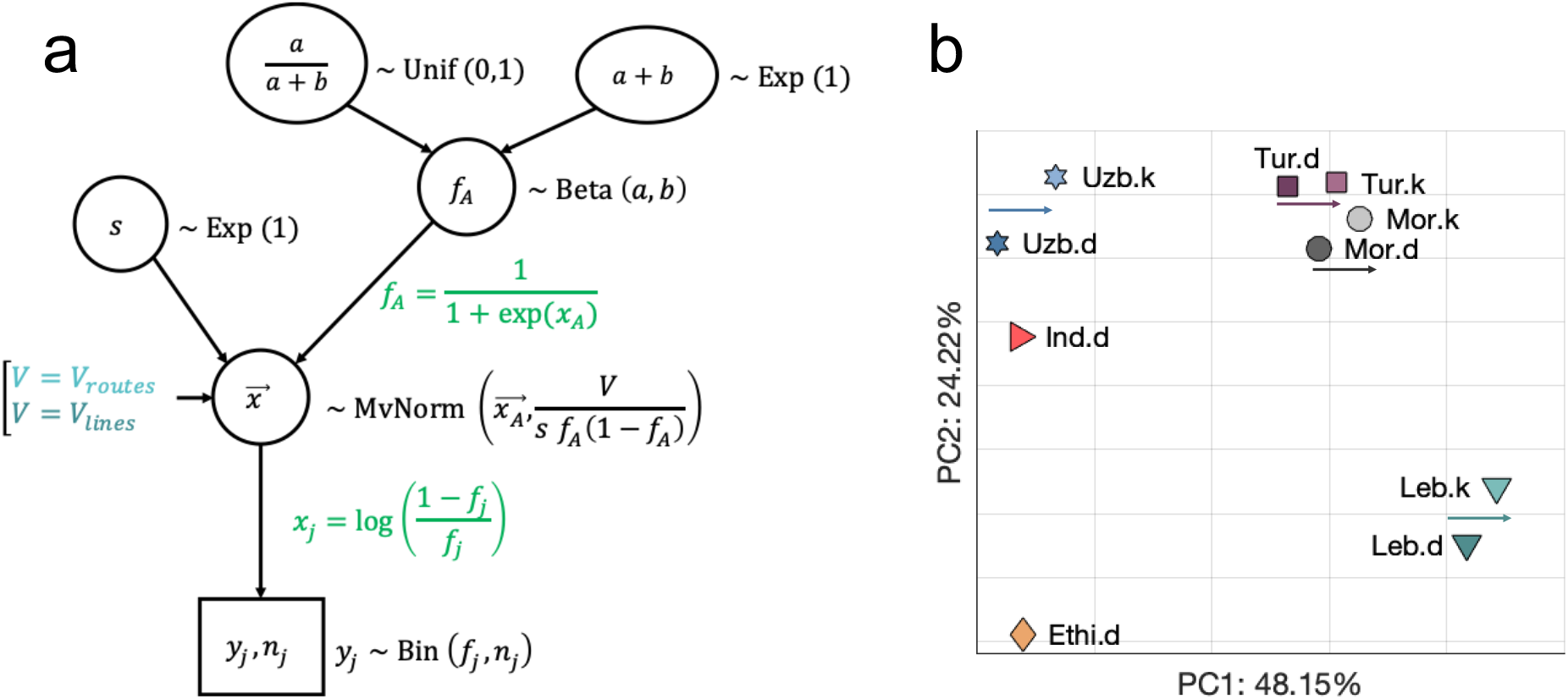
**(a)** Popdisp, the hierarchical Bayesian model describes the spread of chickpea population within each region. We consider that a region consists of *J* sampling locations connecting together by a binary path from the center towards locations. *j*-th location is characterized with *y*_*j*_ allele counts in *n*_*j*_ genotyped variants; *y*_*j*_ and *n*_*j*_ are known values. We assume that *y*_*j*_ is a result of Binomial sampling with *n*_*j*_ trials and *f*_*j*_ probability of success (the allele frequency in the location). Allele frequencies, as fractions or percentages, are constrained (i.e. sum up to 1 or 100%), which requires the transformation of all *f*_*j*_ into *x*_*j*_ being in line with BEDASSLE (Bradburd et al., 2013) and compositional data analysis (CoDA) (Aitchison, 1986; Pawlowsky-Glahn and Buccianti, 2011). The vector 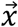 follows the multivariate normal distribution, its mean is the transformed allele frequency in the center, *x*_*A*_, and the covariance matrix is proportional to covariance matrix *V* reflecting the binary path. We tested different paths: constructed under the ‘trade routes’ hypothesis and ‘linear’ hypotheses. Allele frequency in the center has the Beta prior distribution with *α*and *β*parameters. **(b)** PCA plot of allele frequencies estimated under the ‘trade routes’ hypothesis. Arrows represent the shift from desi to kabuli populations within one region.

Chickpea centers of diversity have rich archaeological records, and several domestication scenarios have been proposed based on these. The wild progenitor of *C. arietinum* is *C. reticulatum*, a rare species found in a small area of south-eastern Turkey (Abbo et al., 2003a). Because Turkey (and Syria) also harbor several archaeological sites with the earliest remains of cultivated chickpea (ca 9500 ybp) (Abbo et al., 2003b; Tanno and Willcox, 2006), this region is generally accepted as the origin of chickpea. Based on the archaeological records, chickpea then spread throughout ancient world, reaching western-central Asia (Uzbekistan) and the Indus Valley ca 6000 ybp, the Mediterranean basin (Lebanon, Morocco) ca 5500 ybp, and Ethiopia ca 3500 ybp. While the chickpea migration relationships between Turkey, Lebanon, India and central Asia are supported by archeological records, the exact dispersal and admixture history of chickpea within the Mediterranean Basin and to Ethiopia are anyone’s guess.

The *C. arietinum* L. history gets more complicated due to the presence of two distinct types: ‘desi’ and ‘kabuli’, which differ in size/morphology, color and surface of seeds (Purushothaman et al., 2014) (Figure 1a). Desi and kabuli types have sometimes been designated as subspecies *microsperma* and *macrosperma*, respectively (Moreno and Cubero, 1978), although these older taxonomic terms do not reflect a crossing boundary or substantial molecular genetic differentiation (Varma Penmetsa et al., 2016). The desi type is considered to be ancestral and resembles wild progenitors (*C. reticulatum* and *C. echinospermum*) more than kabuli. It was proposed that kabuli was once selected from the local desis, and then spread; however, the region of origin is not known.

We utilized the genotyped landraces from Vavilov’s collection to test the ambiguities in chickpea history and reconstruct migration routes of both desi and kabuli types in the following way. We first obtained robust estimates of allele frequencies in 10 chickpea populations (6 desis: Turkey, Uzbekistan, India, Lebanon, Morocco, and Ethiopia, and 4 kabulis: Turkey, Uzbekistan, Lebanon, and Morocco). For this purpose, we developed the **popdisp** model (**pop**ulation **disp**ersals), which considers geographical locations of chickpea sampling sites, the nonequal number of samples in locations, and, most crucially, possible ways of chickpea dispersals within a region. We examined two hypothetical dispersals for each of 10 populations and get estimates of allele frequencies in populations’ centers. Then, we used these frequencies to test admixture events in the Ethiopia and Morocco desi chickpea, as well as two different hypotheses about the geographical origin of kabuli varieties and their admixtures with local desis. For these tests, we developed the **migadmi** method (**mig**rations and **admi**xtures), which, instead of existing approaches (TreeMix (Pickrell and Pritchard, 2012) and MixMapper (Lipson et al., 2013)) can cope with more than two source populations and estimate multiple and nested admixture events.

## Results

### Population structure

The chickpea dataset consists of 421 samples (landraces), which can be separated into ten subpopulations based on origin (Turkey, Uzbekistan, India, Lebanon, Morocco, or Ethiopia) and chickpea types (desi and kabuli); there are no kabulis among Ethiopian and Indian landraces in our historical collection (Figure 1a). PCA analysis of samples demonstrated 4 clusters imperfectly correlated with geography, except one cluster with a specific signal to the Ethiopia desis (Figure 1b). The first principal component mostly reflected the difference between desi and kabuli (Figure 1b; see distribution of variance explained in Supplementary File 1). Analysis of the mean pairwise Fst values demonstrated that 10 populations are split into 3 subpopulations reflecting the geographic proximity and overshadowing two chickpea types (Figure 1c): [Turkey-Lebanon-Morocco], [India-Uzbekistan], and Ethiopia. The PCA and Fst results are in line with the previous attempt (Varshney et al., 2019) to decipher the migration and domestication history of chickpea accessions that also revealed region-specific clustering and no clear patterns of desi/kabuli differentiation.

A hierarchical clustering of the landraces based on SNP distance confirmed (Supplementary File 1) that desi-kabuli separation is imperfect, and landraces from different geographical regions are also mixed. To detect unknown population structure we used ADMIXTURE (Alexander et al., 2009), but this did not reveal a clear number of ancestral populations in our dataset (K): the cross-validation error monotonically decreased with no minimum while increasing K from 1 to 20. Similar to the Fst analysis, ADMIXTURE plots for K=3 and K=7 (Supplementary File 1) indicated visually distinct geographic patterns (Turkey-Lebanon-Morocco, India, Uzbekistan, and Ethiopia) but not desi/kabuli separation.

### Chickpea dispersals within geographic regions

Prior to testing migrations and admixtures for 10 chickpea populations: 6 desis (from Lebanon, Morocco, Turkey, Uzbekistan, India, and Ethiopia) and 4 kabulis (from Lebanon, Morocco, Turkey, and Uzbekistan), we estimated allele frequencies in them. Due to the non-uniform distribution of sampling locations in regions and nonequal number of samples in each location, mean allele frequencies in each population can be biased as mean statistics are sensitive to outliers. To get more robust estimates, we developed a model, **popdisp** (Figure 2a), which considers different scenarios for dispersals within a geographic region and takes into account landrace-specific effects. The structure of the model was inspired by BayPass (Gautier, 2015), and processing of allele frequencies was performed as in BEDASSLE (Bradburd et al., 2013) and compositional data analysis (CoDA) (Pawlowsky-Glahn and Buccianti, 2011).

We hypothesized that each region had one trade center, where chickpea was first introduced, and considered two scenarios for subsequent dispersal within the region. In the first scenario, dispersal within each region proceeded by the transport of seeds to local villages via roads and paths. As a result, the genetic relatedness in local landraces would be predicted by the net of regional trade routes. This scenario was contrasted with simple diffusion, so that genetic differences between landraces would be explained by geodesic distance. We called these two scenarios “trade routes” and “linear”, respectively (Figure 2a).

For each region, the center of diffusion was assumed to be the ancient city closest to the geographical mean center for landraces sampled in the region: Axum (Ethiopia), Volubilis (Morocco), Diyarbakir (Turkey), Heliopolis (Lebanon), Ayodhya (India), and Marakanda (Uzbekistan). Then, we constructed two possible contrast binary paths from centers towards sampling sites. The first was estimated using a ‘least-cost’ model, which have emerged as an explanatory framework reflecting transportation routes in archaeology (Figure 1a). The second was constructed using a neighbour-joining algorithm based on linear distance from sampling sites to the center. Differences between paths for regions are shown in Supplementary File 2.

We estimated SNP frequencies in 10 populations under the trade routes and linear scenarios separately and discriminated between them by the Bayes factor (BF, a ratio of the likelihoods). In all cases (except the Lebanon desi population) the “trade route” scenario was strongly favored (Supplementary File 6). Therefore, we concluded that the dispersal from trade centers to farming villages within regions occurred along the ‘trade route’ travel paths and took allele frequency estimates based on this model for further analysis. PCA analysis of the obtained frequencies demonstrated both splitting of populations into geographic subgroups and desi/kabuli differentiation (Figure 2b). Moreover, all kabuli populations are close to their regional desis, but shifted in one direction along the first PC axis. This may reflect a common origin.

### Origin of desi landraces in Morocco and Ethiopia

The desi chickpea type resembles the wild progenitor and is considered ancestral. Its spread between regions is partly known from archaeology: chickpea was domesticated in Turkey and then introduced into India, Uzbekistan and Lebanon. We set these four populations as sources with known phylogeny (black-coloured subtree in Figure 3b). Ethiopian and Moroccan chickpea desi populations appeared later, and their sources are not known (Figure 3a).

**Figure 3.**
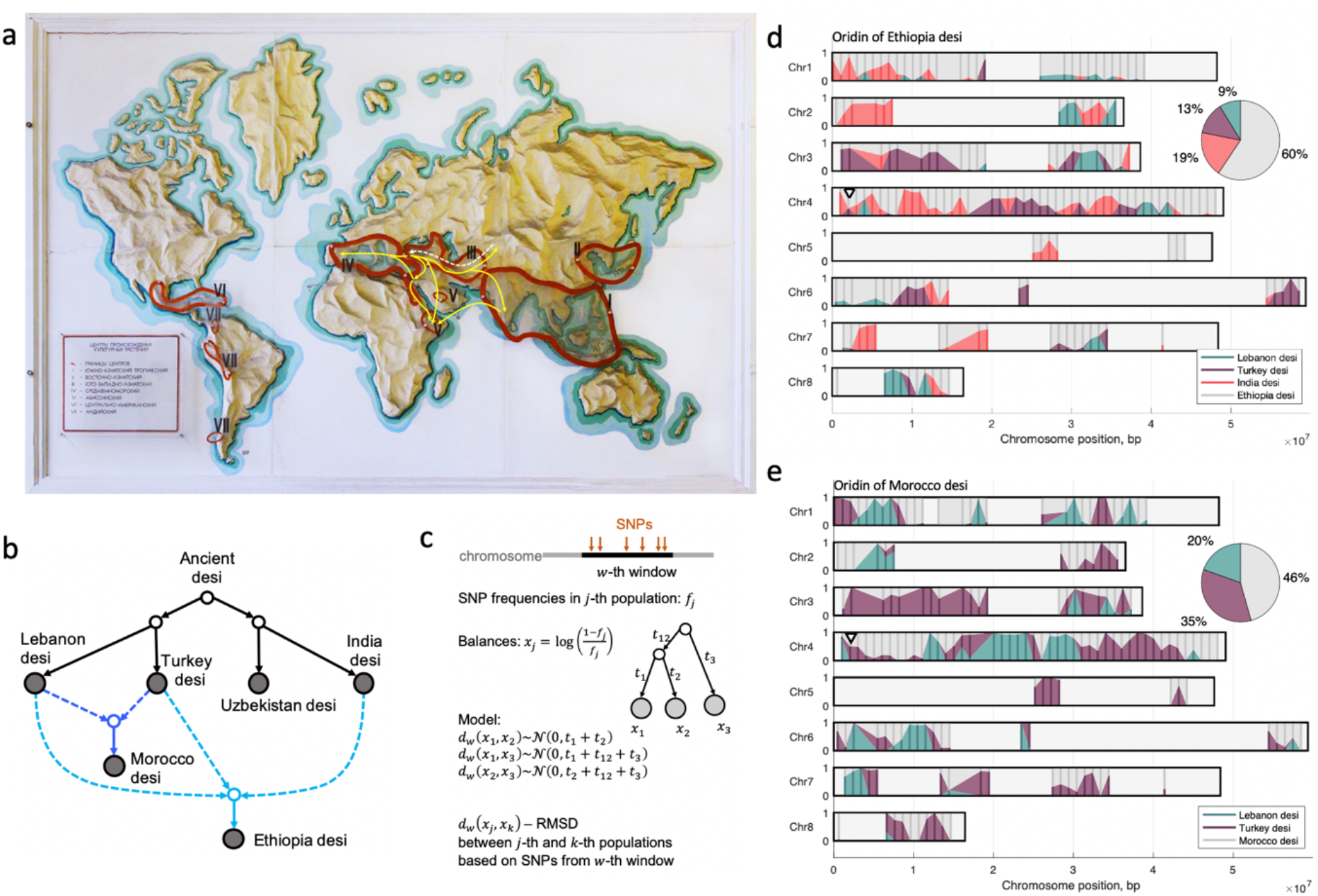
Possible spread of desis between centers of domestication. (A) Vavilov’s centers of domestication (outlined in red) and our hypothesized paths of the desi spread shown as yellow lines (some of which are known and some are tested). The map is from the Vavilov Institute of Plant Genetic Resources (Photo: A. Igolkina). (B) Model of desi’s spread: black lines are known paths of diffusion; we tested the two pathways colored light and dark blue. (C) Parametrization of an admixture event in our model. First, we split each chromosome in a sliding window technique; each *w*-th window is a set of SNPs. Instead of vectors of SNP frequencies for populations, we use vectors balances. We assume that the distance between vectors of balances shortened to the window follows the normal distributions with covariance equal to the corresponding admixture tree’s distance. (D) Distribution of the contribution of Lebanon (green), Turkey (purple), and India (red) ancestral desi populations into Ethiopian desi along chromosomes. (E) Distribution of contribution of Lebanon (green) and Turkey (purple) desi ancestral populations into Moroccan desi along chromosomes.

Two alternative hypotheses exist about the chickpea colonization of Ethiopia. Based on Ethiopian national legend, the Queen of Sheba, a mysterious figure in the Hebrew Bible, is the “founder” of Ethiopia. The Bible tells the story about her visit to Jerusalem (the Gospels of Matthew 12:42, and Luke 11:31), that is in line with Ethiopians highlanders having a clear Semitic connection exemplified by their Semitic language group (Amharic) and genetic similarity with Jewish people (Behar et al., 2010). Based on this, chickpea in Ethiopia might have a Middle Eastern origin. On the other hand, Ethiopian landraces are smaller-seeded and dark-colored, like most Indian varieties. This suggests a South Asian origin of chickpea in Ethiopia. Thus, the genome of these Ethiopian varieties could be admixed with alleles traced back to ancestral populations from Turkey and Lebanon or India. A similar question stands for Moroccan chickpea landraces (Mediterranean Basin), with contributions from either Turkey or Lebanon or both.

Existing methods, like TreeMix (Pickrell and Pritchard, 2012) and MixMapper (Lipson et al., 2013), are not sufficient to test complex historical hypotheses of the chickpea dispersion directly. First, neither of these tools allow both admixed and source populations to diverge after the admixture event. Second, they limit the number of source populations to 2. Third, while TreeMix can estimate multiple admixture events, and MixMapper can cope with two nested admixtures, there is no tool that can do both. Finally, neither tool considers directly the irregularity of admixture traces along the genome, which can be pronounced if the admixture event happened far in the past. We developed a new method, **migadmi** (Figure 3c), which overcomes the above-mentioned limitations. We also applied TreeMix and MixMapper to our dataset and compared their results with ours (Appendix 6).

For Ethiopian desis, the dominant source is India (19%), which has a contribution that is almost as large the cumulative contribution of Lebanon and Turkey desis, 21% (Figure 3d). Thus more than a half of Ethiopian desi’s variance is not represented in ancestral populations, which is in line with the previous analysis, where Ethiopia represents a distinct cluster (Figure 1b,c). These predictions are in agreement with TreeMix results indicating [Turkey-Lebanon] and India origins of Ethiopian desi, while MixMapper suggests that Ethiopian desi is a mixture of desi from Turkey (60%) and India (40%) (Appendix 6). In spite of general agreement of migadmi predictions with TreeMix and MixMapper, we believe that this newly introduced method provides more realistic picture of chickpea colonization in Ethiopia as it takes into account accumulation of individual variances in both mixed and source populations after the admixture event and is able to decompose the variance of mixed population along the chromosomes. Indeed, our analysis demonstrated that non-uniformity of admixture events along chickpea’s chromosomes is strongly pronounced -some regions are admixed by only one source population (e.g. the beginning of chromosome 3 and the middle of chromosome 4 have mainly contribution from Turkish desi population), while other regions have input from several (Figure 3d).

We found that Moroccan desis are derived from both Turkish (35%) and Lebanese (20%) sources (Figure 3e). This result supports the hypothesis of multiple migration routes from West Asia towards Morocco around the Mediterranean Basin. The TreeMix analysis identified Moroccan desi with the Turkish-Lebanese clade (closer to Turkish populations, than Lebanese) with possible India admixture. MixMapper suggested that Moroccan desis are of Turkish origin with an admixture of Lebanese (98%) and Indian (2%) desis (Appendix 6). As the Indian desi influence on Moroccan desi is small, we concluded again that migadmi predictions of a Moroccan origin generally agree with predictions of TreeMix and MixMapper but provide additional information about admixture traces along the chromosomes.

### Origin of kabuli chickpea

The origin of kabuli domestication is unknown. Based on linguistic evidence, one may hypothesize that kabulis arose in Central Asia, and are named after Kabul city (in modern Afghanistan). On the other hand, it is logical to suggest that kabulis arose in West Asia (modern Turkey) but later than desis, as kabulis are distributed in regions neighboring to Turkey and have long been thought to be modern introductions to India and Ethiopia(van der Maesen, 1984). Mulitiple geographic origins are possible. Although desis and kabulis have much in common, modern breeding programs generally keep them separate, likely due to differences in adaptive requirements and market preferences (Purushothaman et al., 2014; Roorkiwal et al., 2014; Varshney et al., 2019).

### Origin of kabuli chickpea

The origin of kabuli domestication is unknown. Based on linguistic evidence, one may hypothesize that kabulis arose in Central Asia, and are named after Kabul city (in modern Afghanistan). On the other hand, it is logical to suggest that kabulis arose in West Asia (modern Turkey) but later than desis, as kabulis are distributed in regions neighboring to Turkey and have long been thought to be modern introductions to India and Ethiopia (van der Maesen, 1984). Although desis and kabulis have much in common, modern breeding programs generally keep them separate, likely due to differences in adaptive requirements and market preferences (Purushothaman et al., 2014; Roorkiwal et al., 2014; Varshney et al., 2019). Because desi type is considered to be more primitive and ancestral it is not unreasonable to assume that kabuli’s spread between centers of secondary diversification had an influence from local desis. However, both kabuli’s origin and migration history with possible desi influences remain unclear.

To identify the origin of kabulis, we draw alternative admixture graphs of population relatedness. The first assumes the dispersal of kabuli chickpea from Turkey’s Fertile Crescent (Figure 4a) and the second reflects a Central Asian origin (modern Uzbekistan) with subsequent movement back to Turkey (Figure 4b). Parameters for the black-coloured part of the graphs in Figure 4a,b were taken from the previous analysis of desi populations, the remaining parameters were estimated with the migadmi model. The optimal likelihood of the former graph is higher, but not significantly. Therefore, to determine the kabuli’s origin, we analysed fractions of variance in each mixed population explained by its sources.

**Figure 4.**
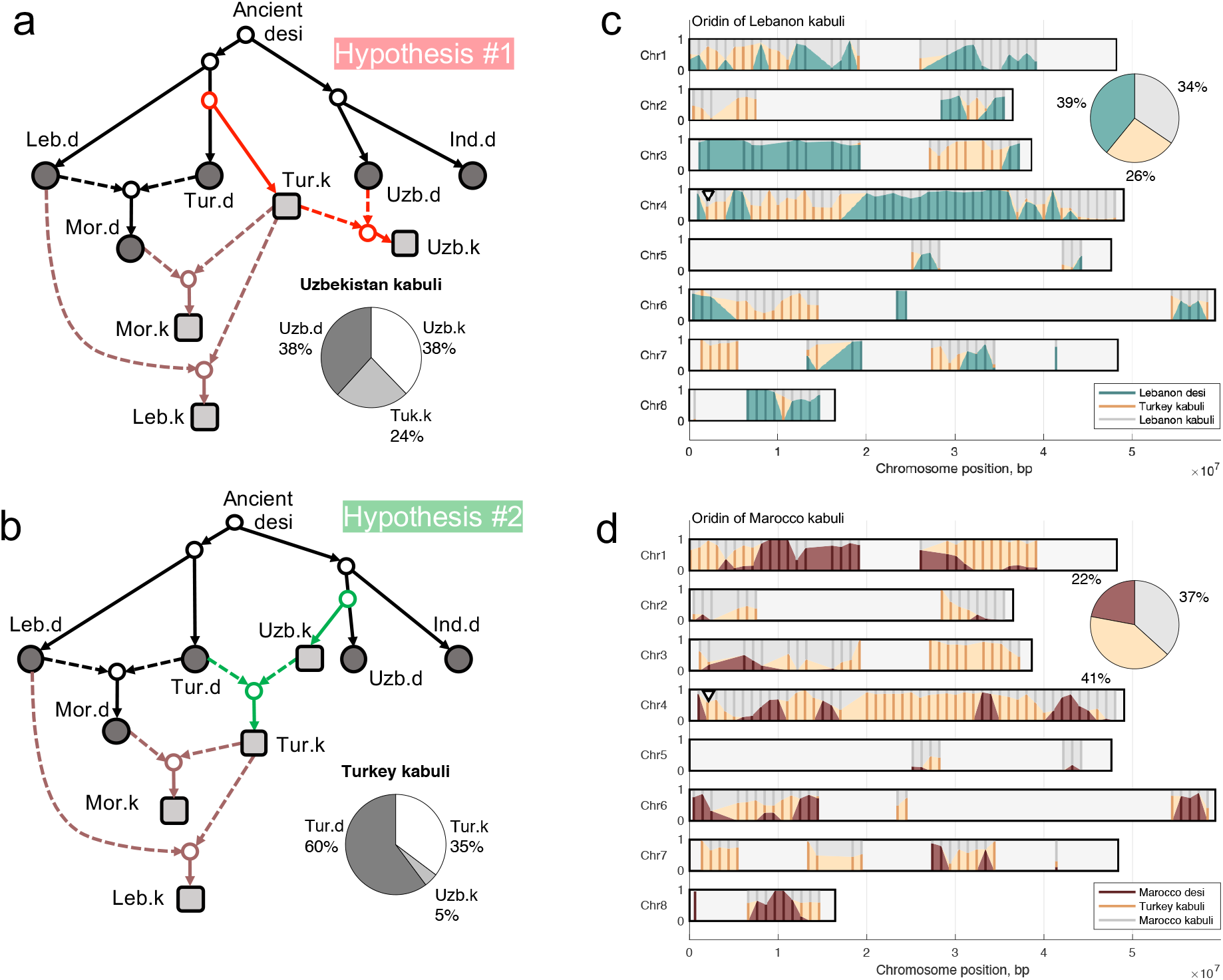
Analysis of the origin of kabuli chickpeas. (a) Paths of kabuli movement assuming that they originated in Turkey. The pie plot reflects the decompositions of Uzbekistan kabuli variance. (b) Paths of kabuli movement assuming that they originated in Uzbekistan (Kabul). The pie plot reflects the decompositions of Turkish kabuli variance. (c) Decomposition of the Lebanon kabuli origin along the chromosomes. (d) Decomposition of the Moroccan kabuli origin along the chromosomes. Triangle marks chromosomal regions associated with kabuli.

Under the Central Asian assumption of kabuli origin, the influence of Uzbeki kabuli on Turkish kabuli is very small (5%), while, under the Turkey origin hypothesis, the influence of Turkish kabuli on Uzbeki kabuli was about 5 times larger (24%) (pie plots in Figures 5a,b). The larger contribution of assumed source to a kabili population indicates Turkey as the likely origin of kabuli. The analysis of PCA plot (Figure 2c) demonstrated the shift of all kabuli populations along the first PC axis, and the direction of this shift is not “towards Uzbekistan.” TreeMix analysis did not reveal significant patterns of kabuli admixture, while the MixMapper indicated the same pattern as we found (Appendix 6). Overall, we do not observe support for a kabuli origin in Central Asia with introgression back to Fertile Crescent populations, and we thus cautiously conclude that kabuli originated in the Turkish region.

**Figure 5.**
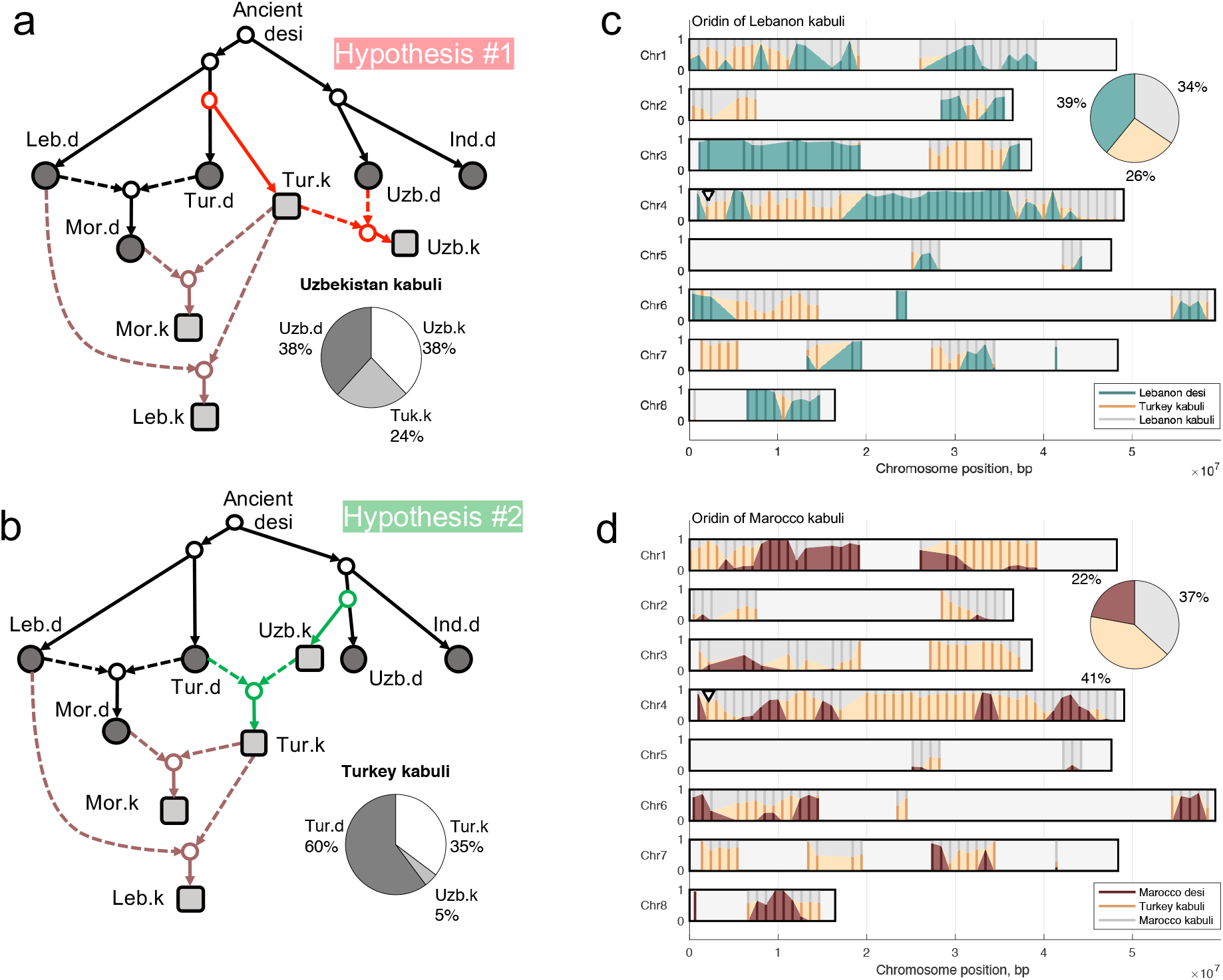
Analysis of the origin of kabuli chickpeas. (a) Paths of kabuli movement assuming that they originated in Turkey. The pie plot reflects the decompositions of Uzbekistan kabuli variance. (b) Paths of kabuli movement assuming that they originated in Uzbekistan (Kabul). The pie plot reflects the decompositions of Turkish kabuli variance. (c) Decomposition of the Lebanon kabuli origin along the chromosomes. (d) Decomposition of the Moroccan kabuli origin along the chromosomes. Triangle marks chromosomal regions associated with kabuli.

Moroccan and Lebanese kabuli varieties appear to be highly related to both local desi and Turkish kabuli (pie plots in Figures 5c,d). The proportion of Turkish admixture in the Moroccan kabuli population (41%) is higher than in the corresponding desi populations (22%), evidence that the desi landraces spread earlier than kabuli landraces, and have had more time to diverge and accumulate their own variance. The mixed origin of Moroccan and Lebanese Kabulis was also demonstrated by MixMapper, but the influence of local desi was higher than the influence of Turkish kabuli in both cases (60%) (Appendix 6). Analysis of regions admixed by Turkish kabuli in Moroccan and Lebanese kabuli chromosomes reveals common patterns (Figures 4c,d) and highlights the chromosomal regions associated with kabuli (Appendix 7). For example, the beginning of the fourth chromosome, which contains markers for chickpea flower color, the basic difference between desi and kabuli varieties (marked as triangle on the Figures 2d,e and 3c,e) (Varma Penmetsa et al., 2016) contains clear introgression from the Turkish kabuli ancestral population. Of note, that chromosomal region in Ethiopia appears to be derived from India (Figure 2d).

## Conclusion

We have tested chickpea migration and admixture hypotheses directly, by formulating dispersal scenarios (Figures 3 and 4) based on historical evidence. We observed that the Ethiopian desi population was derived not solely from the Fertile Crescent, but almost equally from India and the Fertile Crescent (Turkey-Lebanon). Likewise, a uniform variation pattern around Mediterranean (Varshney et al., 2019) has been clarified into two likely land routes of migration from the Fertile Crescent, via Sothern Europe and North Africa.

Another question which we addressed was the origin of kabuli, the light-colored chickpea type, which presumably originated from a local desi population. According to the analysis we performed this region is Turkey. We observed no evidence for kabuli’s Central Asia origin and spreading back to the Fertile Crescent as was speculated previously (Varshney et al., 2019).

To test the migration and admixture hypotheses, we developed two methods. The first model is popdisp, which estimates allele frequencies in the population, under the assumption of a particular dispersal model within the region. We considered two reasonable physical agents of migration: traders or diffusion that approximates continuous-time stochastic process. Our assertion was that genomic resemblance between accessions can reflect either ‘least-cost path’ trade route distance between sample sites or linear distance between them. Our analyses unambiguously favour the former hypothesis (Figure 1a). In the future it will be interesting to apply this approach to species with different dispersal strategies, for instance comparing crops like round-seeded chickpea to human-associated weeds like spiky-podded *Medicago* capable of long-distance transport with livestock ^24^ or wind dispersed species. For the latter, we would expect distributions to track wind currents only, with no resulting signature of dispersal along historic trade routes.

The second model is migadmi, which estimates multiple and nested admixture hypotheses with more than two sources and demonstrates the admixture patterns along the chromosomes. Both models describe changes in allele frequencies in line with Wright-Fisher drift model and utilize logit transformation as in BEDASSLE (Bradburd et al., 2013) and compositional data analysis (CoDA), the most appropriate framework for working with frequencies, fractions, percentages and ratios. This approach allows to easily extend migadmi to work with not only biallelic SNPs, but also with multiallelic sites or haploblocks.

## Materials and methods

### Dataset

The chickpea dataset (Cicer arietinum L.) consists of 421 accessions from the Vavilov Institute of Plant Genetic Resources (VIR) seed bank. These accessions were genotyped by sequencing (GBS), and 56,855 segregating single nucleotide polymorphisms (SNPs) were identified. These SNPs were further filtered to meet requirements for minor allele frequency (MAF) >3% and genotype call-rate >90%. 2,579 SNPs in 421 accessions passed all filtering criteria and were retained for further analysis (Sokolkova et al., 2020).

### Spatial Data and Distance Calculations

To estimate physical distances between sample locations of chickpea accessions, we took into account the spherical model of Earth and geodesic measurements. We used the Projection Wizard web application (Šavrič et al., 2016) to select an accurate projection for regions with locations onto the two-dimensional surface (Appendix 1).

To calculate distances between pairs of locations, we used the Least-cost path model (Douglas, 1994) (instead of pure geodesic measurements), the explanatory framework for the movement of goods in archeology. This approach calculates the least “cost” distance of a path, that can be interpreted as an amount of time or energy that it would have taken to travel along the path. This approach is useful in the absence of historical data on exact movement routes, and it takes into account the change in elevation, the hiking function (which is used in archeological and ethnographic applications (Gorenflo and Gale, 1990), geo-climatic Holocene data, and a mask of water bodies (see detailed description in Appendix 1).

For each of six regions (Ethiopia, Morocco, Turkey, Lebanon, India, and Uzbekistan), we estimated possible locations of chickpea diffusion centers combining current knowledge of World Centers of Diversity and historical data for locations of ancient cities that were prominent trading centers during ancient times. Using the spatial statistics tools, we calculated the mean center for each region and then compared the centers’ locations with known ancient trade/cultural centers (Ancient World Mapping Center. University of North Carolina, Chapel Hill, http://awmc.unc.edu/awmc/map_data/shapefiles/strabo_data/). As a result, we selected the following historic settlements closest to the mean centers: Axum (Ethiopia), Volubilis (Morocco), Diyarbakir (Turkey), Heliopolis (Lebanon), Ayodhya (India), and Marakanda (Uzbekistan).

### Model for diversification within clusters

The model describing populations dispersals is implemented in Python package popdisp (https://github.com/iganna/popdisp).

### Model

We developed popdisp, a Bayesian hierarchical model (Figure 2a) that describes historical diversification of chickpea populations within a geographical region. We hypothesize that each geographic region contains *M* populations originated from one center (ancestral population) and spread towards *M* locations. Each population is composed of individuals genotyped for *N* unlinked (independent) biallelic SNPs; the missing data is possible and does not require the imputation. We pooled the data from all individuals in a population; for *j*-th population and *i*-th SNP, we defined the total counts of non-reference (alternative) allele –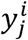– and the total count of all variants at this SNP –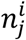. Values 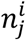 are not the same across all SNPs in *j*-th population due to the missing data. We assume that frequency of the alternative allele for *i*-th SNP in *j*-th population is 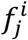, and the observed 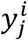 follows the Binomial distribution: 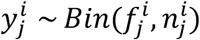.

Within a region, we modelled population spread along a given binary-branching path from the ancestral population, which is characterized by respective frequency 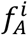. We assumed that population allele frequencies change under the genetic drift in line with the Wright-Fisher model and theory of Compositional Data analysis (CoDA). The CoDA theory states that frequencies (as well as percentages or fractions) are meaningless when considered alone, as they sum up to one, hence, the only balances between frequencies do make sense. According to the CoDA, we applied the isometric log-ratio (ilr) transformation to allele frequencies, and, in case of biallelic SNPs, it is the logit transformation as used in BEDASSLE (Bradburd et al., 2013):

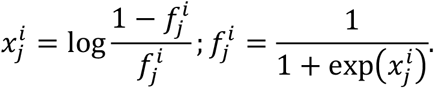

New variable 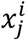 means the log-balance between frequencies of reference and alternative alleles, and is not bounded, i.e., can take values in (−∞, +∞). The latter allows us to model correlations between population frequencies using Multivariate normal distributions without artificial truncation, which is necessary when the model operates with non-transformed frequencies(Gautier, 2015).

To describe the genetic drift of allele frequencies along the binary-branching paths, we modified the approach proposed in TreeMix(Pickrell and Pritchard, 2012) and BayPass(Gautier, 2015). In the Wright Fisher model, the expected value and variance of allele frequency in *j*-th population are 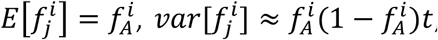, where *t* is the amount of genetic drift, which has occurred along the path from the ancestral population to *j*-th population. To match these first two moments after ilr-transformation of allele frequencies (Appendix 2), the following should be satisfied: 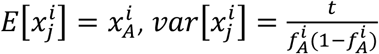.

Using the logic of model construction from TreeMix(Pickrell and Pritchard, 2012) and Gaussian model for changing log-balances, we get that 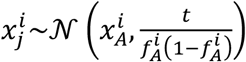, where *t* is proportional to the cumulative path from the ancestral population to *j*-th population. Using the Felsenstein’s approach (Felsenstein, 1973), we model the change of log-balances along the binary-branching path with Multivariate normal distribution:

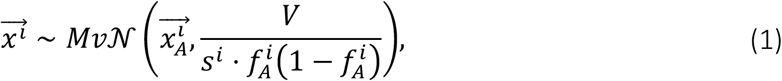

where 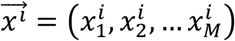 is the constant of proportionality specific for *i*-th SNP, *V* is *M*× *M* matrix, which reflects the covariance structure between *M*population based on the binary-branching path. This path can be represented as a binary tree structure with ancestral population at the root and *M* leaves (Figure 2b). On the diagonal, matrix *V* contains cumulative branch lengths from the tree root to respective leaves, and the off-diagonal elements are equal to sum of common branches for respective pair of populations(Felsenstein, 1973). We compute values in *V* matrix based on known length of binary-branching path and scale it, so that the mean value of diagonal elements should equal to one.

### Prior probabilities and MCMC

For each SNP, model has the following parameters: the allele frequency in the ancestral population, log-balances of allele frequencies for *M* populations, and the constant of proportionality. To get estimates, we constructed Bayesian model with the following prior distributions for parameters.

For 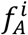, we proposed uninformative beta prior, *Beta(a*^*i*^, *b*^*i*^) with uniform prior for the mean, 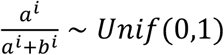, and exponential prior for the so-called “sample size”, *s*^*i*^*∼Exp*(1). We also assume the exponential prior for constant of proportionality: *s*^*i*^ ∼ *Exp*(1).

The complexity of the model does not allow the use of Gibbs Sampling. Instead, we performed the algebraic inference of derivatives for log posterior distribution and run Hamiltonian Monte Carlo sampling algorithm (Neal, 2012) in pyhmc (https://pythonhosted.org/pyhmc/) to get parameter estimates. For each chickpea subpopulation we ran 3 MCMC chains of length 50,000 and traced the Gelman-Rubin convergence diagnostic (<1.1) and effective sample size.

To conclude which model of chickpea dispersal within a region is more probable, we separately got estimates on *V* matrix calculated for trade routes and linear distances. Then we compared log posterior values between two estimates (Supplementary File 6).

### Model for migration between clusters

The migadmi model describing **mig**rations and **admi**xtures of populations is implemented in Python package **migadmi** (https://github.com/iganna/migadmi).

To test hypothetical migration routes of chickpea between regions, we created a model based on the same assumptions as used in the model for population spread within a region. We consider *P* populations characterised with vectors of log-balances of allele frequencies, which are obtained from the previous analysis. We denote log-balances of allele frequencies of *i*-th SNP in *j*-th populations with 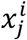.

A migration hypothesis is set by the binary tree, which branch lengths are parameters. Based on the migration hypothesis, we construct the parametrized covariance matrix *V* and matrix *D* containing variances of differences between log-balances: *D*_*jk*_ = *V*_*jj*_ + *V*_*kk*_ − 2*V*_*jk*_. Then, we can construct the following likelihood function (Appendix 3):

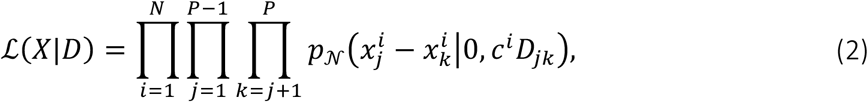

where *N* is a number of SNPs, *X* is the matrix of log-balances for all SNPs and all populations *c*^*i*^ is a SNP-specific scale parameter.

The likelihood (2) contains a unique scale parameter, *c*^*i*^, for each SNPs, making the model overparametrized. To reduce the number of parameters, we applied the sliding window technique. We divided each chromosome into overlapping windows of the same size almost equal to the LD, 3 · 10^8^ bp; the step parameter in the sliding window was 1 · 10^8^. As the density of SNPs along chromosomes is not uniform (Supplementary File 5), windows contained different numbers of SNPs; those with less than 10 SNPs were filtered out.

We assumed that SNPs within each window are probably linked and had evolved with a similar rate. This assumption allows us to avoid *c*^*i*^ parameters (set it to 1), and infer objective function proportional to log-likelihood (see Appendix 4):

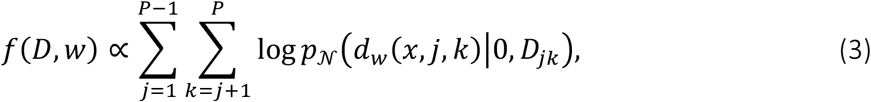

where *d*_*w*_(*x, j, k*) is a root mean square distance between *j*-th and *k*-th populations, computed on SNPs from *w*-th window (see Appendix 4), log *p*_𝒩_ denotes the log-density of normal distribution. We estimate parameters in *D* matrix separately for each window.

### Modeling admixture events

We developed a new model of admixtures which considers that (i) admixture events happened long ago and all populations (both source and mixed) accumulated their own variance after the event, (ii) number of source populations in one event are not constrained, i.e., can be higher than 2, (iii) several admixture events can be analyzed simultaneously, and (iv) admixtures can form a hierarchy, i.e., a mixed population in one admixture event can be a source in another event.

Let population *y* be a mixture of *Q* sources 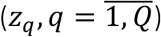, which are precursors of *Q* current populations 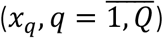. We parametrized this admixture event with the following variables: *t*_*y*_ – own variance of the mixed population; *w*_*q*_ – weights of source populations, 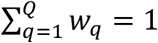 *α*∈ [0,1] – part of own variance of *x*_*q*_ which is common with *z*_*q*_ (see Appendix 5). To avoid overparameterization, we set the regularization on *w*_*q*_ with the Dirichlet prior (all concentration parameters, λ, equal to 0.9).

To test an admixture hypothesis, we (i) constructed the corresponding tree with admixture events, (ii) parametrized *V* and *D* matrices based on the tree, (iii) estimated parameters maximizing the objective function (4).

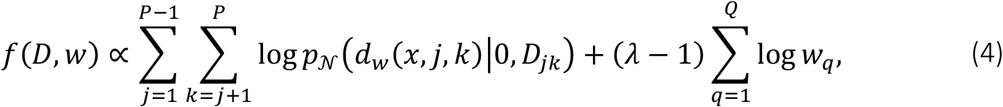

## Appendix 1 Geographic distances between locations

### Projection

The map projection used to represent a geographic region on a flat surface plays a critical role when measuring distances (such as distances between regions), areas or assessing shape or direction. Whenever a spherical model of Earth is projected onto two-dimensional surface, distortions of one or another kind are introduced, altering these variables to a different degree. Our project area stretches from the Iberian Peninsula through the Mediterranean Ocean, swinging south to Ethiopia and further covering parts of Central Asia, to the West India, laying below 60 degrees North to the Equator. That spatial extent and the ultimate focus on extracting physical distances, called for Equidistant Conic Secant projection, which is characterized by having two standard parallels (as opposed to Tangent projections that have only one standard parallel). This projection has proved practical since Classical times (Snyder, 1993). We used the Projection Wizard web application (Šavrič et al., 2016) to select accurate angular and linear parameters for the transformation.

### Calculation of Distances

It is typical to use geodesic measurements of distance between pairs of points in landscape genomics (Abebe et al., 2015) and although these can yield adequate results, they do not take full advantage of genomic data to provide insights into historical patterns of trade and diffusion. Least-cost path models (Douglas, 1994) have emerged as an explanatory framework for movement of goods in archeology (Kantner, 1997). This approach of calculating the distance of a path with the least “cost” (interpreted usually as change in elevation) provides a mechanism, in the absence of historical data on exact movement routes, to estimate the time and energy that it would have taken to travel from location to location. Pairwise distances between concentrations of accessions were calculated both using geodesics as is typical in landscape genomics (Abebe et al., 2015) and as least-cost paths with slope and water bodies defining landscape friction, following a trend to use three-dimensional spatial modeling to predict trade routes between ancient settlements (Herzog, 2014; van Lanen et al., 2015). We used the hiking function, which has been used in archaeological and ethnographic applications (Gorenflo and Gale, 1990) to assign resistance along with a cost surface accounting for climatic conditions.

We created a cost surface using selected geo-climatic Holocene data sets, mask of water bodies, and weighted elevation gradient, rescaled to a common scale. We used the following climatic layers: maximum temperature of the warmest month, minimum temperature of the coldest month and precipitation of wettest month for past conditions (Mid-Holocene), obtained from WorldClim, Version 1.4 database, MIROC-ESM GCM (Hijmans et al., 2005). Temperature and Precipitation ranges were ranked in accordance with ASHRAE Thermal Comfort chart (Hoyt et al., 2013).

A slope layer was created from the world elevation (GTOPO30) and reclassified according to the Tobler function (Tobler, 1993). In addition, a water mask was created to mask out water bodies. We then used Weighted Overlay tool of ArcGIS to create a cost surface layer, where each pixel had a value of the least accumulative cost distance from or to a source of interest. Supplementary File 7 describes scheme of classification for each layer and its relative weight in building cost surface.

One hypothesis is that movement between sites always goes through historical centers of trade before dispersing out to rural villages. In this exploratory analysis we converted least-cost paths between mean centers that could have served as the foci of crop dispersion, using data acquired from the Ancient World Mapping Center, UNC GIS, into vector format and construct a road network for the whole area.

The cost distance layer was further used to prototype paths between cities (regional centers of dispersion) as well as within each cluster. The resultant least-cost path rasters were converted to vector format, cleaned of duplicates and served as base data for building a road network. We then employed ArcGIS Network Analyst functionality to build a road network that encountered for terrain relief and point connectivity, and to retrieve distance values between and within spatial clusters. Straight-line geodesic distances were calculated with the ESRI ArcGIS Near tool.

### Selection of Centers of Diversification

We estimated the number and locations for hypothetical centers of diffusion by combining current knowledge of regions that served as World Centers of Diversity (Corinto, 2014), cluster analysis of our accessions’ locations, and historical data for locations of ancient cities that were prominent trading centers during ancient times (Ancient World Mapping Center, n.d.).

We applied ArcGIS clustering analysis and spatial statistics tools to group all accessions into six clusters based on geographic locations and spatial constraints, and to calculate mean center for each cluster. We then compared the locations of the mean centers with known ancient trade / cultural centers (Ancient World Mapping Center, n.d.) and selected a historic settlement closest to each calculated mean center: Axum (Ethiopia), Volubilis (Morocco), Diyarbakir (Turkey), Heliopolis (Lebanon), Ayodhya (India), and Marakanda (Uzbekistan)

## Appendix 2 First two moments of ilr-transformed allele frequencies

Let a population be described by the frequency of alternative allele of a biallelic SNP, *f*. The population comes out from the ancestral one with the allele frequency *f*_*A*_ under the Wright Fisher model of genetic drift. In the Wright Fisher model, expected value and variance of allele frequency are 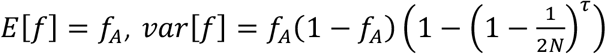, where *τ* is the number of generations separating current and ancestral populations, and *N* is the size of diploid population. Using the Binomial approximation, 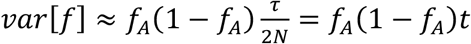, where *t* can be considered as the amount of genetic drift.

We applied the ilr-transformation for allele frequencies and obtained 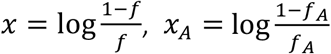. These new variables mean the log-balance between reference and alternative allele frequencies in the current and ancestral populations. Using Taylor expansions, the second order approximation of the expected value of *x* is *x*_*A*_, and the approximation of variance is the following:

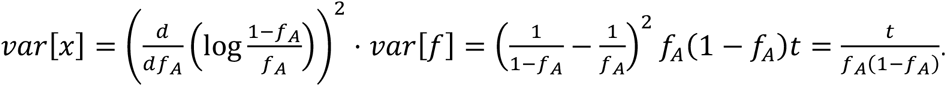

## Appendix 3 Estimates for branch parameters of a tree

Let’s consider *P* populations originated from one ancestral state and a binary tree depicting their migration history; all tree branch lengths are parameters. Each population is characterized by log-balance of allele frequencies for a SNP, *x*_*i*_. In the model for population spread within a region, it has been assumed that 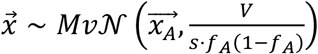, where 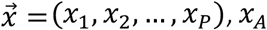, *x*_*A*_ is the log-balance of allele frequency in the root of the tree (ancestral state). However, in testing historical hypotheses, there is no given information about the ancestral state: *f*_*A*_ is not known, position of the root in the binary tree is parametrized. Therefore, it is impractical to include *f*_*A*_ into the model and use the above-mentioned multivariate normal distribution.

To avoid the use of *f*_*A*_, we propose an approach which considers total variance between populations instead of covariance. Let covariance matrix between populations, *V*be obtained based on the fully parametrized binary tree according to Felsenstein’s method(Felsenstein, 1973) (see Example on Figure A1). Then, we can obtain a matrix *D*, which elements are proportional to variances of the difference between log-balances:

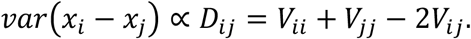

Based on Gaussian changing log-balances, we get: (*x*_*i*_ − *x*_*j*_) ∼ 𝒩 (0,*c* · *D*_*ij*_), where *c* is a constant of proportionality covering 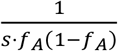.

To get maximum likelihood estimates of the tree branch length based one SNP, the following likelihood function can be written:

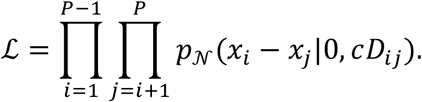

**Figure A1.**
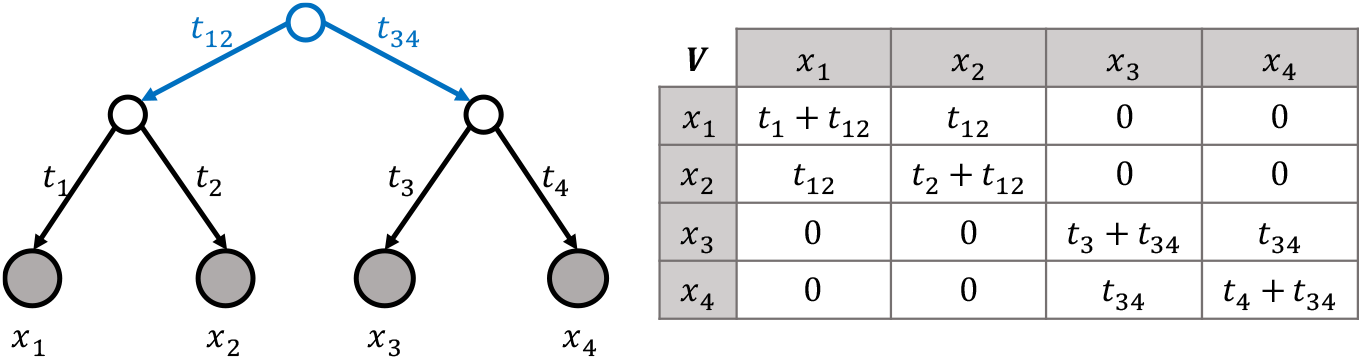
Example of constructing matrix V based on the tree with parametrized branches.

## Appendix 4 Inference of likelihood function for a set of linked SNPs

A “window” is a segment on a chromosome of length equal to a predefined value (≈LD) that contains a subset of SNPs. We assumed, that, within each window, SNPs are probably linked and they had evolved with a similar rate. Let *G*_*w*_ be a set (group) of SNPs corresponding to *w*-th window, and *s*^*w*^ be a scale, specific for this window and reflecting the rate. For *i*-th SNP in *j*-th population, we denote log-balances of allele frequency with 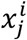. Then, the Likelihood function for log-balances of allele frequencies in the *w*-th window is:

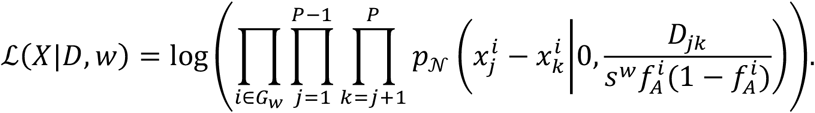

where 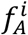 is the allele frequency of the ancestral state. This value is not a parameter, is not known, and plays the scale role. In line with CoDA, we estimate it as 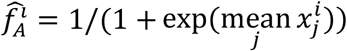. Let denote constant 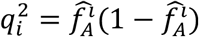, then the likelihood is proportional to:

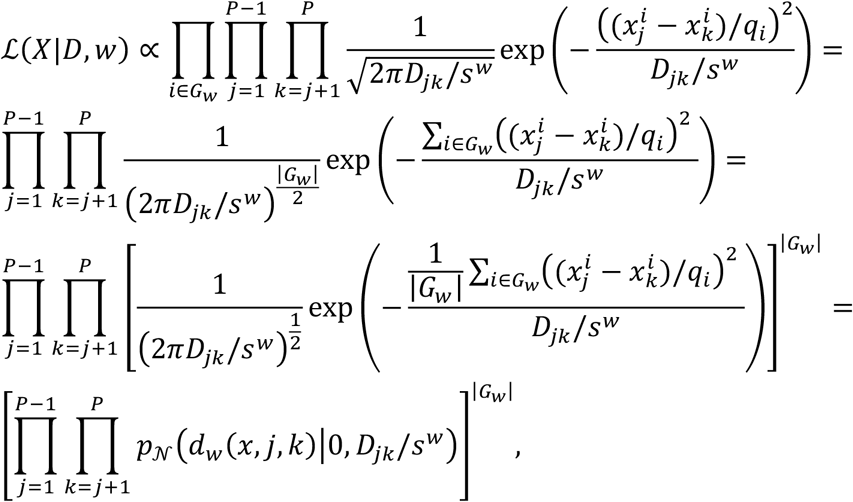

where 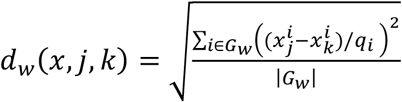 is the normalized root mean square distance between *j*-th and *k*-th populations, computed on SNPs from *w*-th window. However, as matrix *D* is fully parametrized, we can set *s*^*w*^ = 1 without loss of generality. To get parameters estimated, we can remove the power and maximize the following log-likelihood function:

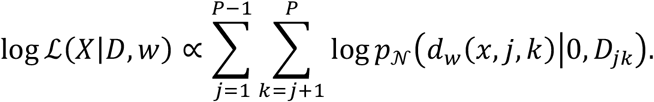

## Appendix 5 Identification of parameters in the mixture model

Consider six populations originated from one ancestral state, and a tree depicting the history of the populations (Figure A2a); *x*_*j*_ is a normal random variable reflecting the log-balance of frequencies for the SNP in population *j* (Figure A2a). We denote lengths of tree branches with *t*_*i*_.

Let the seventh population (having *y* log-balance of frequencies for the SNP) originate by a mixture event of three populations (precursors of *x*_1_, *x*_3_, and *x*_6_), and then evolve independently along the branch with the length *t*_*y*_ (Figure A2b). We assume that the mixture event happened long ago, so that current populations *x*_*i*_ have their own evolutionary history, independent from the sources *z*_*i*_. To carefully consider the mixture event, we introduced weight parameters *w*_*i*_, α_*i*_, β_*i*_, as demonstrated in Figure A2b,e. In our example, the number of additional parameters is 10, and the number of constraints is 4; hence, the number of free parameters is 6. The number of cells in the matrix *D*, which contain additional parameters, is 6, so all free parameters are identifiable in this example. However, in the extreme situation, when all six initial populations can be considered as sources of the mixed one, the number of free parameters reaches 12, and some of them become non-identifiable.

In general, when the initial tree connects *n*_*pop*_ populations and all of them can be sources of a mixed one, the number of free parameters is 2*n*_*pop*_ and number of cells in the matrix *D*, which contain additional parameters, is *n*_*pop*_. Therefore, to avoid this overparameterization we introduce several constraints. First, we assume that all α^*i*^ are equal to each other, and this assumption reduce the number of free parameters to (*n*_*pop*_ + 1) (Figure A2c). Second, we set the regularization on *w*_*i*_ weights using the Dirichlet prior with all concentration parameters equal to 0.9: 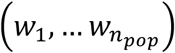 ∼ Dirichlet(0.9 … 0.9). Imitating absorbing states in the genetic drift, this prior tends to pull some weights to zeros, i.e. to put 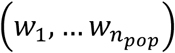 vector closer to the border of *n*_*pop*_-dimensional simplex. These two introduced restrictions make all free parameters in the model identifiable.

**Figure A2.**
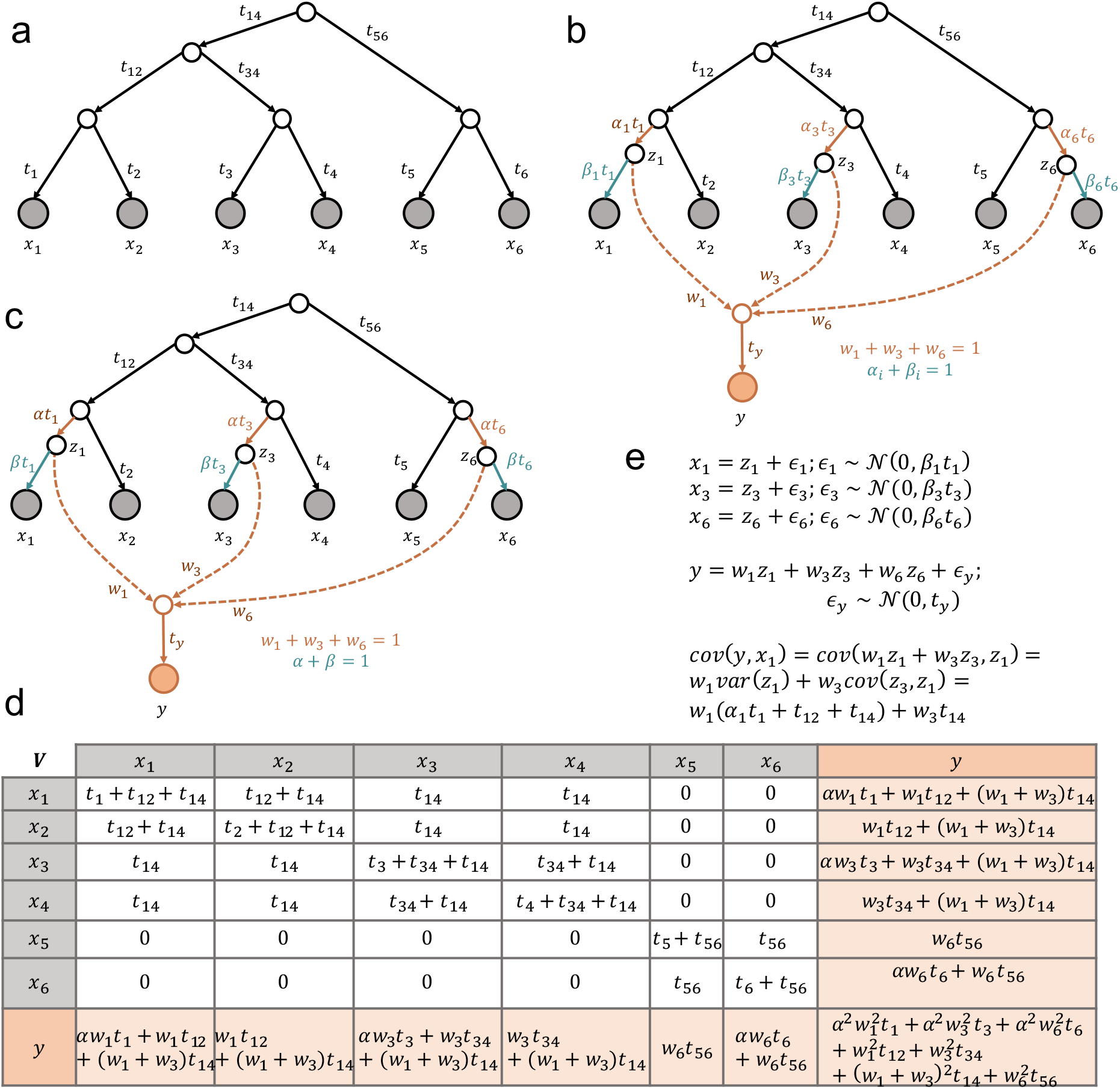
An example tree describing the evolutionary history of 7 populations with admixture; *x*_*i*_ represent the frequency balance of a SNP for *i*-th population, *y* is the population formed with an admixture, *t*_*i*_ are the length of a tree branch, *w*_*i*_, α_*i*_, and β_*i*_ are a weight parameters. The *V*-table demonstrates the variance-covariance matrix *V*for all populations after re-parametrization.

## Appendix 6 Comparison of migadmi results with TreeMix and MixMapper

**Table A1.** Comparison of admixture methods

|  | migadmi | TreeMix | MixMapper |
| --- | --- | --- | --- |
| Admixture of >2 sources | + | - | - |
| Several non-nested admixtures | + | + | - |
| Number of nested admixtures | $\geq 2$ | 0 | 2 |
| Adding admixture event to core tree | + | - | + |
| Admixture pattern along the chromosome | + | - | - |
| Can take the tree as input | + | + | - |
| Accounting for own evolutionary history for both mixed population and source populations | + | - | - |
| Modeling frequencies | Compositional data analysis | Normality assumption | Normality assumption |

To estimate the migration and admixture events in our study, we developed a new method, **migadmi**, because of the limitations of the existing ones, TreeMix (Pickrell and Pritchard, 2012) and MixMapper (Lipson et al., 2013). We created a list of characteristics to compare the packages and found that our method covers and outperforms capabilities of TreeMix and MixMapper: our package copes with estimating multiple complex admixture events with more than 2 sources and demonstrates the admixture patterns along the chromosomes. Moreover, it has two additional features that were not accounted for in previous models.

The first feature is that, migadmi allows populations to get their own variance after admixture events. In the existing approaches, it is assumed that the composite population is a weighted sum of some source populations, and weights sum to 1. However, in reality, almost no population is settled as a net sum of two or more. Ordinarily, when a part of one population appears in a new place, it evolves some period of time getting its own variability, and then if the admixture event happens, the mixed population continues to evolve. As a result, the variance in the admixed population can be factored into contributions from source populations and self-accumulated variance. The latter is especially important if the admixture events happened long ago (e.g., as in our study). Things get more complicated when considering that source populations have also evolved. To avoid modeling the mixed populations as a weighted sum of source ones, we parametrized the own variance of each population after the admixture event.

The second important feature of **migadmi** is the use of ilr-transformed allele frequency instead of allele frequency itself. Allele frequencies, as fractions or percentages, are constrained (i.e. sum up to 1 or 100%), which makes standard statistical methods inapplicable. For example, frequencies cannot be modelled as normally distributed random variables, as the domain of the normal distribution is (−∞, +∞), not [0, 1]. Another problem is presence of negative bias in covariance estimates between frequencies (Aitchison, 1986). Moreover, frequency of one allele is inextricably linked with frequencies of others as they sum to 1. Therefore, modeling frequency changes of one allele cannot be considered without modeling changes in other alleles. To correctly work with frequencies, the theory of compositional data analysis and Aitchison geometry were first established in the end of previous century (Aitchison, 1986)(Pawlowsky-Glahn and Buccianti, 2011). Following this theory, one can independently analyze (*D* − 1) balances between frequencies, instead of *D* frequencies. In case of biallelic SNPs, the balance is the logarithm of the ratio between reference and alternative alleles, and this balance takes values in (−∞, +∞). We adapted the use of balances to model changes of allele frequencies in line with the Wright-Fisher drift model. The balance-based approach was used in both **popdisp** and **migadmi** models.

The direct comparison of migadmi results with TreeMix and MixMapper results is not possible because we used migadmi to estimate complex admixture graphs, which TreeMix and MixMapper cannot cope with (Table A1). However, we performed the standard TreeMix and MixMapper analyses and traced the common and different trends in results.

First, we applied TreeMix and set to estimate 4 events within 10 populations. We used TreeMix in two modes: without tree root specification and with specificationof Ethiopia desi population as a root, the most distinct one (Figure A3). We also used the bootstrap with the size of 35, that equals to the mean number of SNPs in our sliding window technique. Both obtained admixture graphs demonstrated two expectable distant clades in trees: Uzbekistan-India and Turkey-Lebanon-Morocco. However, the obtained trees also contained deviations from the expectations. In the root-specified tree, the Ethiopian desi population is the source for Turkish desi that contradicts the conventional story of chickpea spread (Figure A3a). The root-unspecified tree contains India’s influence on Moroccan desi, which is also unlikely, because these populations are the most distant to each other (Figure A3b).

On the other hand, TreeMix graphs partly support the hypothetical origin of Ethiopian and Moroccan desis. The location of Ethiopian desi on the root-unspecified tree demonstrated its sources from both main clades, which is in line with the mixed origin of this population. In the root-unspecified tree, the Moroccan desi population is located between Turkish and Lebanese populations, while in the root-specified tree, it locates close to Turkey with an admixture from Lebanese desi. Therefore, we may conclude that Moroccan desi is an indirect mixture of Turkish desi and Lebanese desi.

The origin of kabulis is impossible to infer from this tree, however, the root-specified tree indicates that the Uzbeki kabuli has an admixture from the Turkey-Morocco clade, that is in line with our hypothesis, that Uzbeki kabuli is not the source of other kabulis.

**Figure A3.**
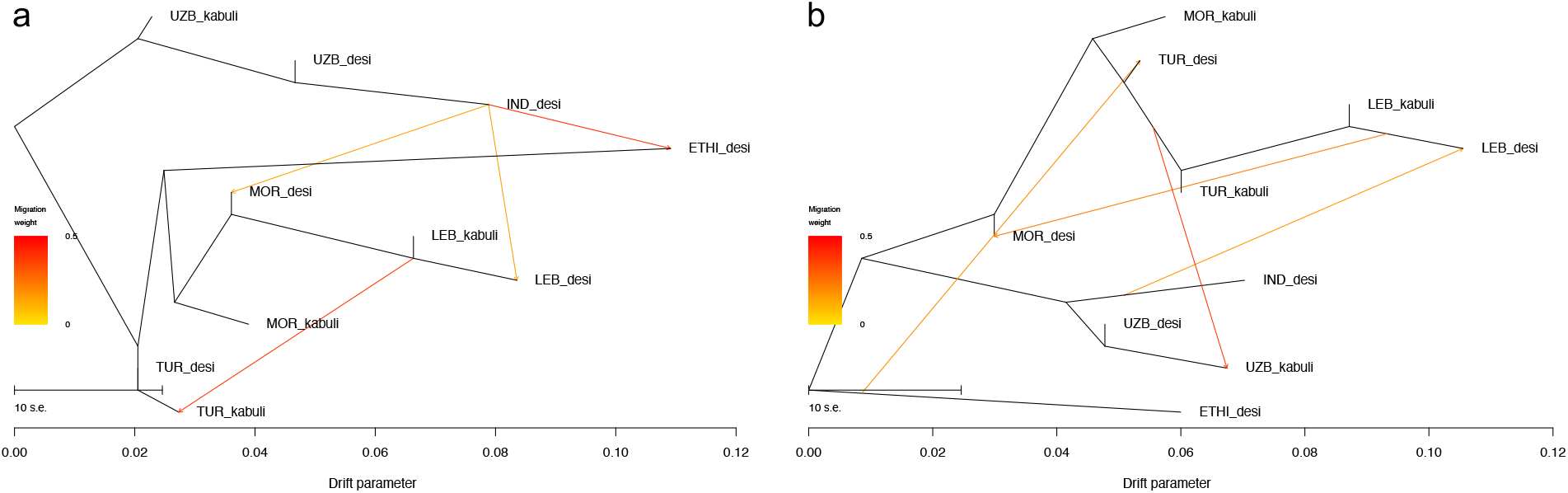
Admixture graphs obtained with the TreeMix package for (a) unrooted tree and (b) rooted with the Ethiopian desi population. Firstly, TreeMix estimates the tree based on all input populations (black branches), and then it introduces admixture events (colored arrows). Color of lines reflects the weight of the admixture from 0 to 0.5.

MixMapper takes source populations as input, then creates a tree on them and tests a mixed population adding it to the tree. We applied MixMapper in the bootstrap mode to match windows from our analysis. We analyzed the origin of Ethiopian desi, taking Turkish, Lebanese, Indian, Uzbeki desis as source populations. MixMapper revealed two sources of Ethiopian desi: Turkish desi (60%) and Indian desi (40%). The direct analysis of Moroccan desi as a mixture from Turkish, Lebanese, Indian, Uzbeki desis revealed that it is as a mixture from Lebanese desi (98%) and Indian desi (2%).

To test the origin of kabuli, we tested two models and compared the admixture coefficients. In the first model, we assumed that Turkish, Lebanese, Indian, Uzbeki desis, and Turkish kabuli are five source populations, and Uzbeki kabuli is a mixture. The direct analysis revealed that Uzbeki kabuli has 62% from Uzbeki desi and 38% from Turkish kabuli. In the second model, we assumed that Turkish, Lebanese, Indian, Uzbeki desis, and Uzbeki kabuli are five source populations, and Turkish kabuli is a mixture. In this case, we found that Turkish kabuli is a mixture of Turkish desi and Lebanese kabuli, so that not from Uzbeki kabuli. Therefore, we may conclude that origin of kabuli is likely Turkey.

Then, we took Turkish, Lebanese, Indian, Uzbeki desis, and Uzbeki kabuli and tested them as sources for Lebanese kabuli and Moroccan kabuli separately. Lebanese kabuli is predicted to be a mixture local desi (60,2%) and Turkish kabuli (30,8%). The Moroccan kabuli was tested in the nested model (as Moroccan desi is also the mixture), which revealed Moroccan kabuli as a mixture of Moroccan desi (60,3%) and Turkish kabul (39,7%).

## Appendix 7 Chromosomal regions associated with kabuli/desi difference

The most pronounced difference between desi and kabuli chickpea types is the flower color. In legumes, this trait is Mendelian and controlled by the so-called A gene (Hellens et al., 2010). For *Pisum sativum* and *Medicago truncatula*, the sequences of this gene can be found at GenBank accessions: GU132940 (MtbHLH) and GU132941 (PsbHLH). We took these sequences, performed the tBLASTn search against Cicer ariethinum genes, and found the match with basic helix-loop-helix protein A located at LOC101506726 locus (2149255-2158629bp, the beginning of chromosome 4).

To verify that this region is associated with desi/kabuli difference, we performed GWAS analysis on the binary trait (belonging to desi or kabuli) using rrBLUP. We found one significant SNP which is located very close to the found homologous LOC101506726 locus. Therefore, we suppose that this locus can be considered as a marker locus for kabuli.

**Figure A4.**
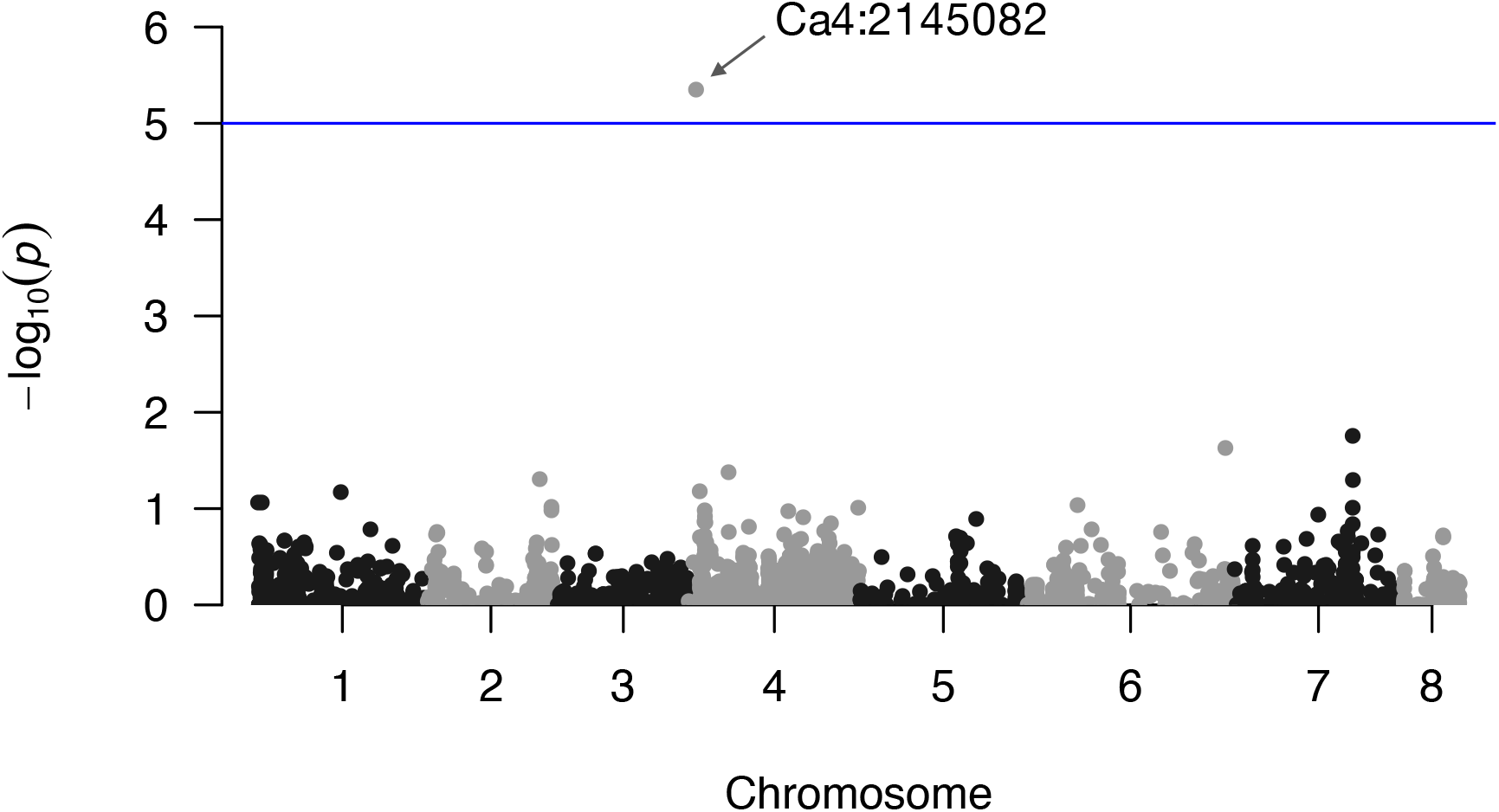
Manhattan plot for GWAS of desi/kabuli binary trait.

## Data Availability

All Illumina data are available from the National Center for Biotechnology database under BioProject PRJNA388691. Processed initial data for the analysis is uploaded to GitHub repositories with the code.

## Code Availability

Code for the popdisp and migadmi analysis frameworks are available at: https://github.com/iganna/popdisp and https://github.com/iganna/migadmi.

## Acknowledgements

The research was supported by RFBR grant 18-29-13033 to A.A.I., M.G.S. and S.V.N.; by a cooperative agreement from Dthe United States Agency for International Development under the Feed the Future Program AID-OAA-A-14–00008 to S.V.N., E.J.B.v.W.; and a grant from the James H. Zumberge Faculty Research and Innovation Fund to S.V.N. and T.L. E.J.B.v.W. is further supported by the USDA Hatch program through the Vermont State Agricultural Experimental Station.

We would like to thank Magnus Nordborg for discussions and his helpful advice on the paper structure.

---

**Supplementary Figure 1.**
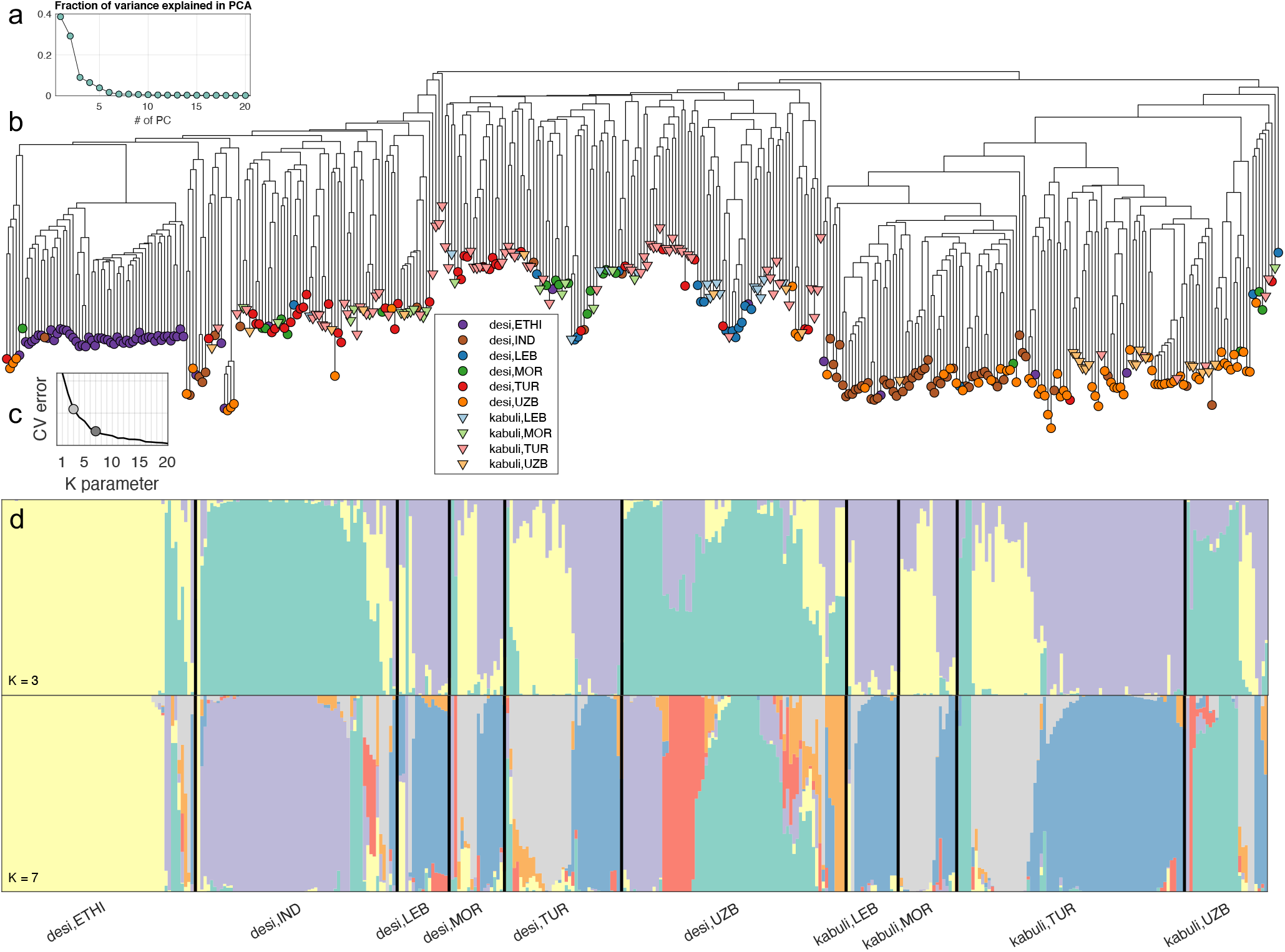
Population structure of chickpea landraces. (a) proportion of variance explained by PCs in PCA analysis on all SNP data (b) Neighbor-joining tree of chickpea accessions using SNP-distance. The ten chickpea subpopulations are marked with different colors. (c) Cross-validation plot for different numbers of ancestral populations used in the ADMIXTURE program. The curve does not show a minimum, that is a criterion for K choice. Two points reflect cross-validation errors for runs demonstrated below. (d) Population structure inferred by ADMIXTURE analysis for K=3 and K=7. Each chickpea sample is represented by a stacked column with K components corresponding to estimated ancestral populations colored differently (components sum to 100%). Samples are ordered according to the ten chickpea subpopulations.

**Supplementary Figure 2.**
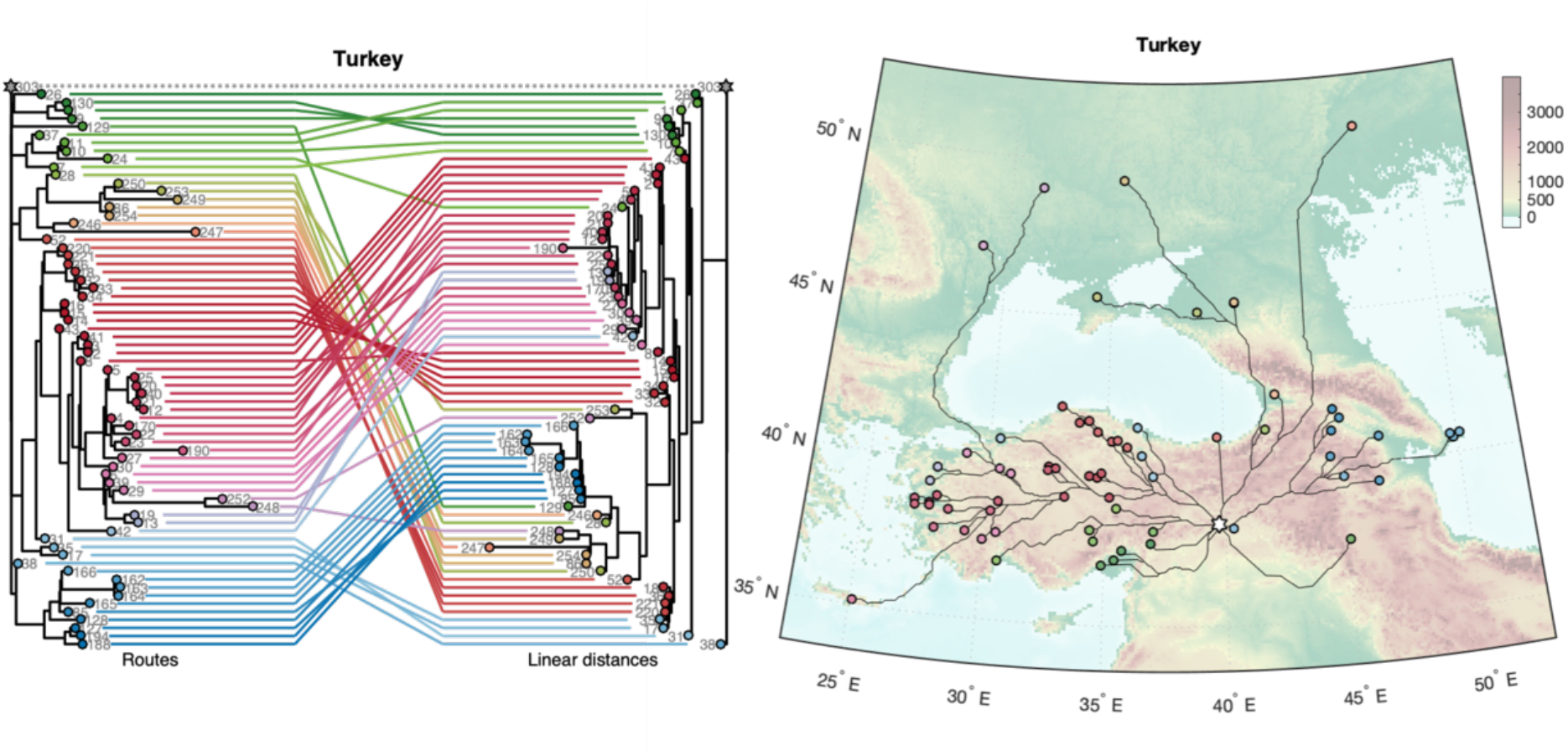
Tanglegram for correspondence between routes and linear distances within the Turkey cluster. Routes of the Turkey cluster on Map; star denotes the center of the cluster.

**Supplementary Figure 3.**
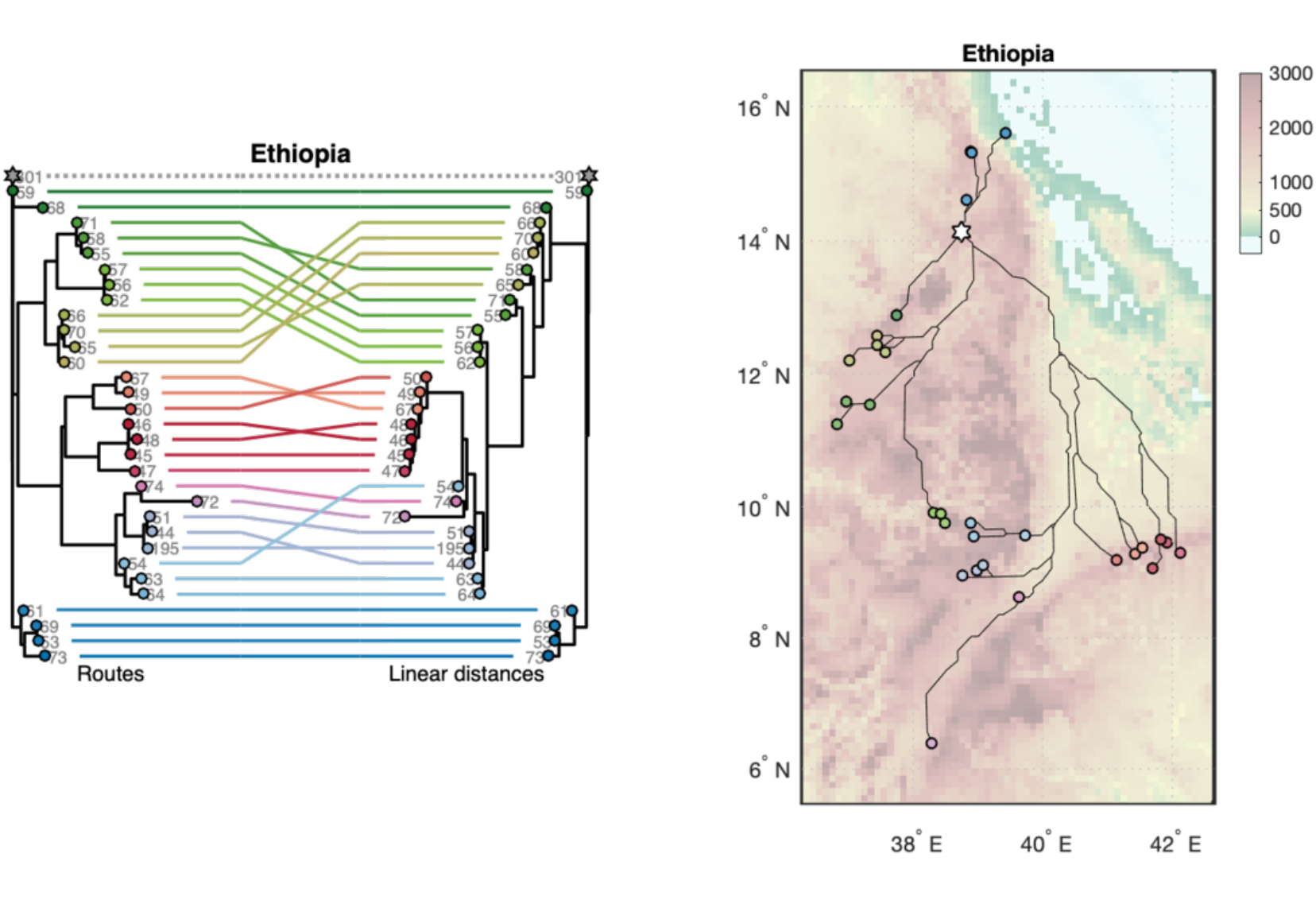
Tanglegram for correspondence between routes and linear distances within the Ethiopia cluster. Routes of the Ethiopia cluster on Map; star denotes the center of the cluster.

**Supplementary Figure 4.**
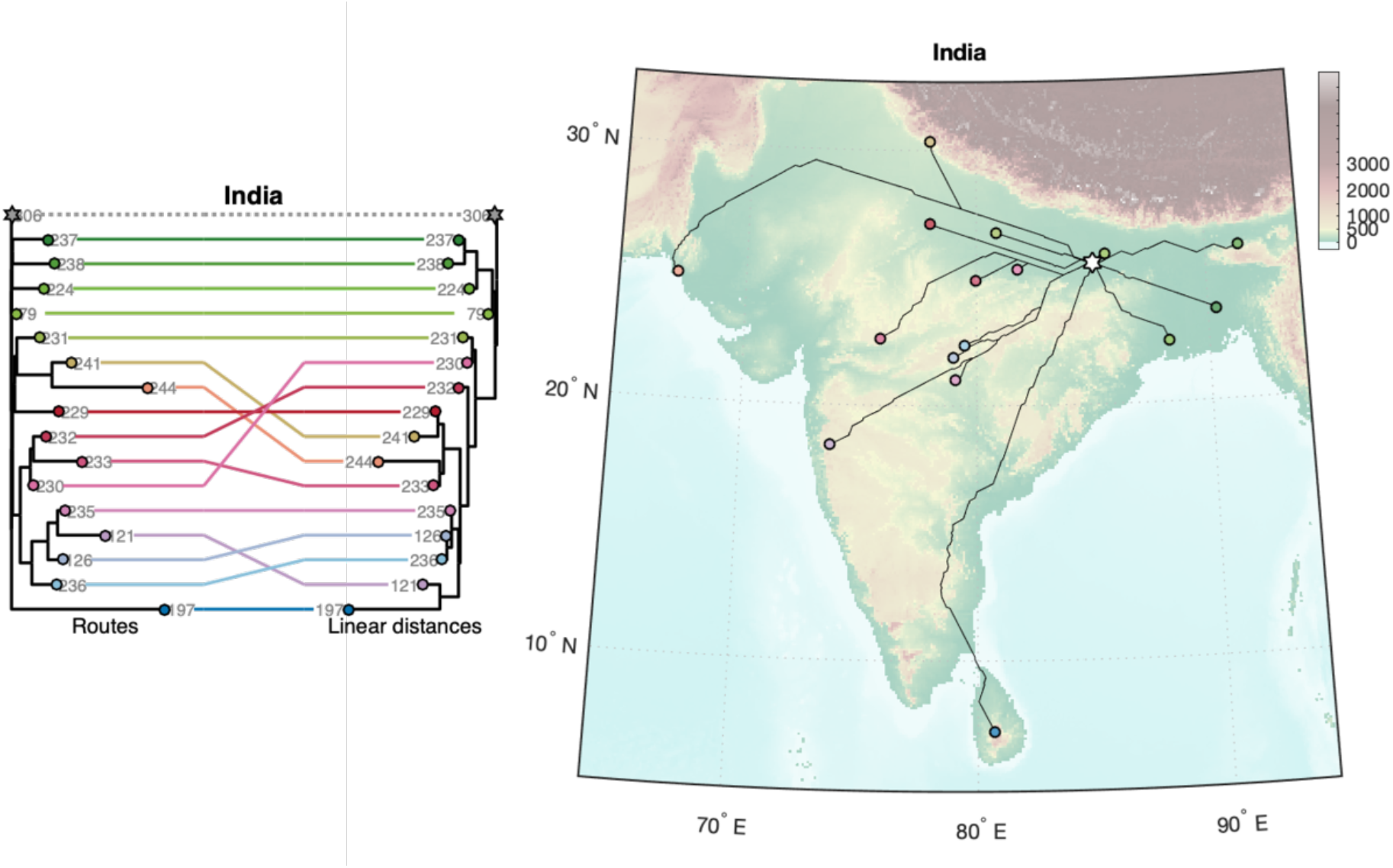
Tanglegram for correspondence between routes and linear distances within the India cluster. Routes of the India cluster on Map; star denotes the center of the cluster.

**Supplementary Figure 5.**
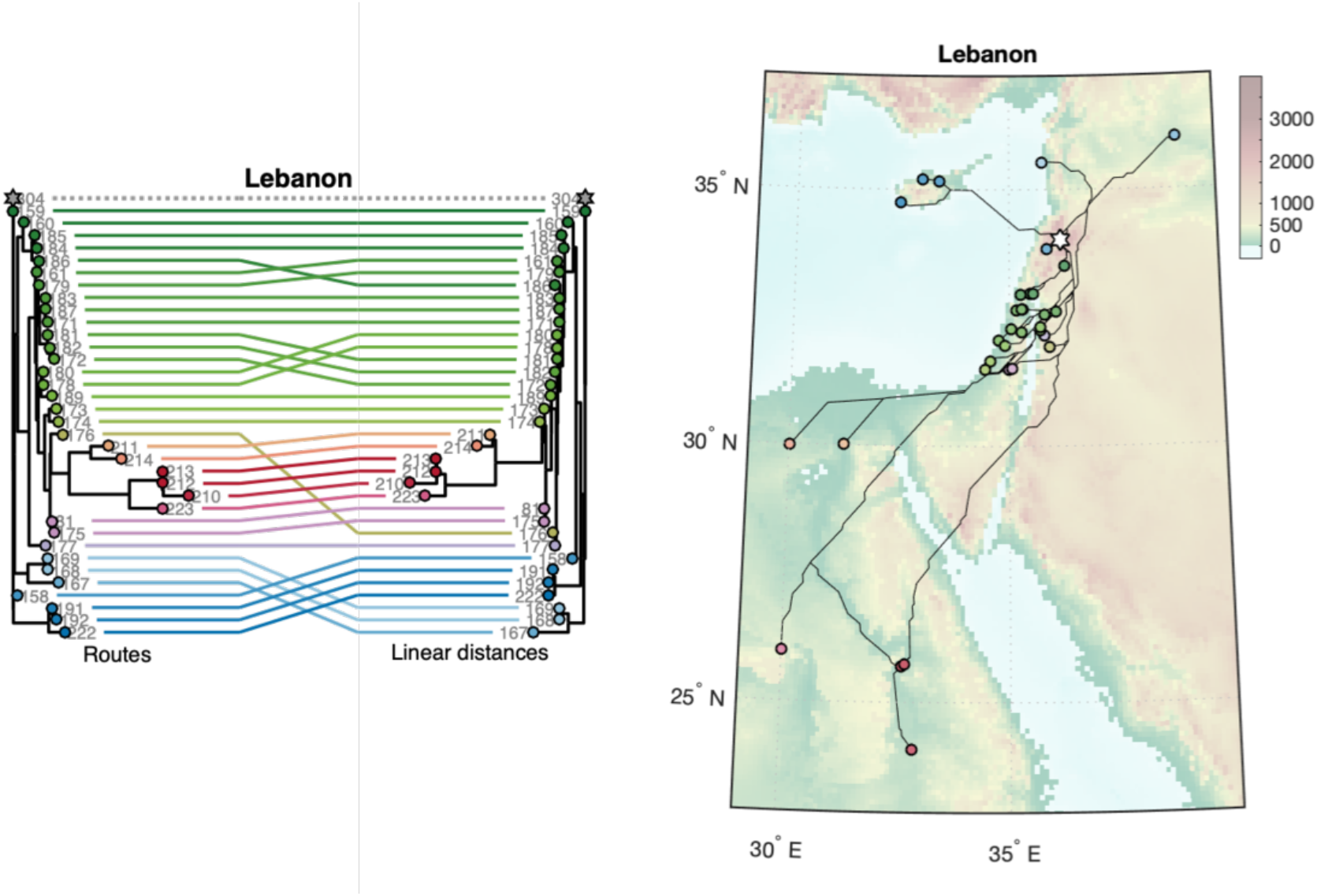
Tanglegram for correspondence between routes and linear distances within the Lebanon cluster. Routes of the Lebanon cluster on Map; star denotes the center of the cluster.

**Supplementary Figure 6.**
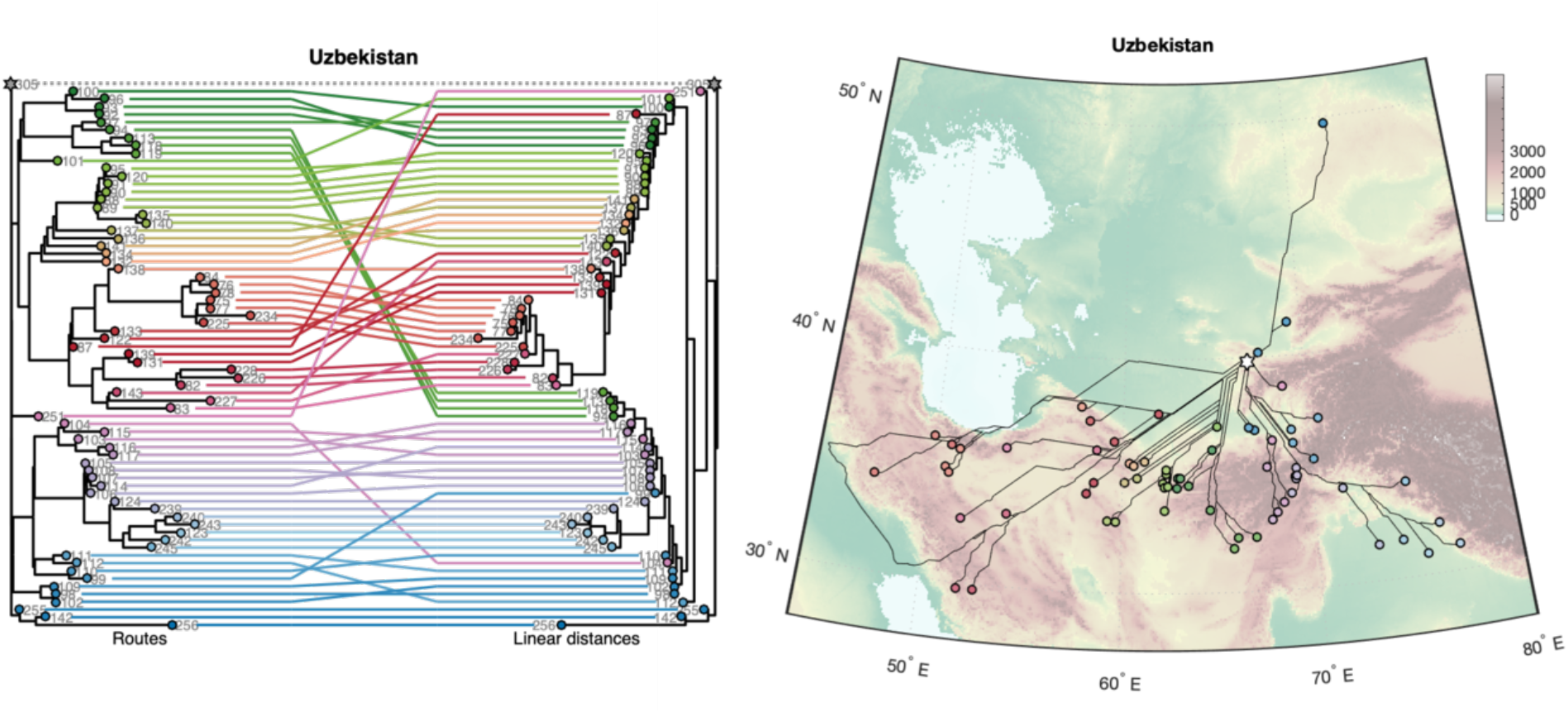
Tanglegram for correspondence between routes and linear distances within the Uzbekistan cluster. Routes of the Uzbekistan cluster on Map; star denotes the center of the cluster.

**Supplementary Figure 7.**
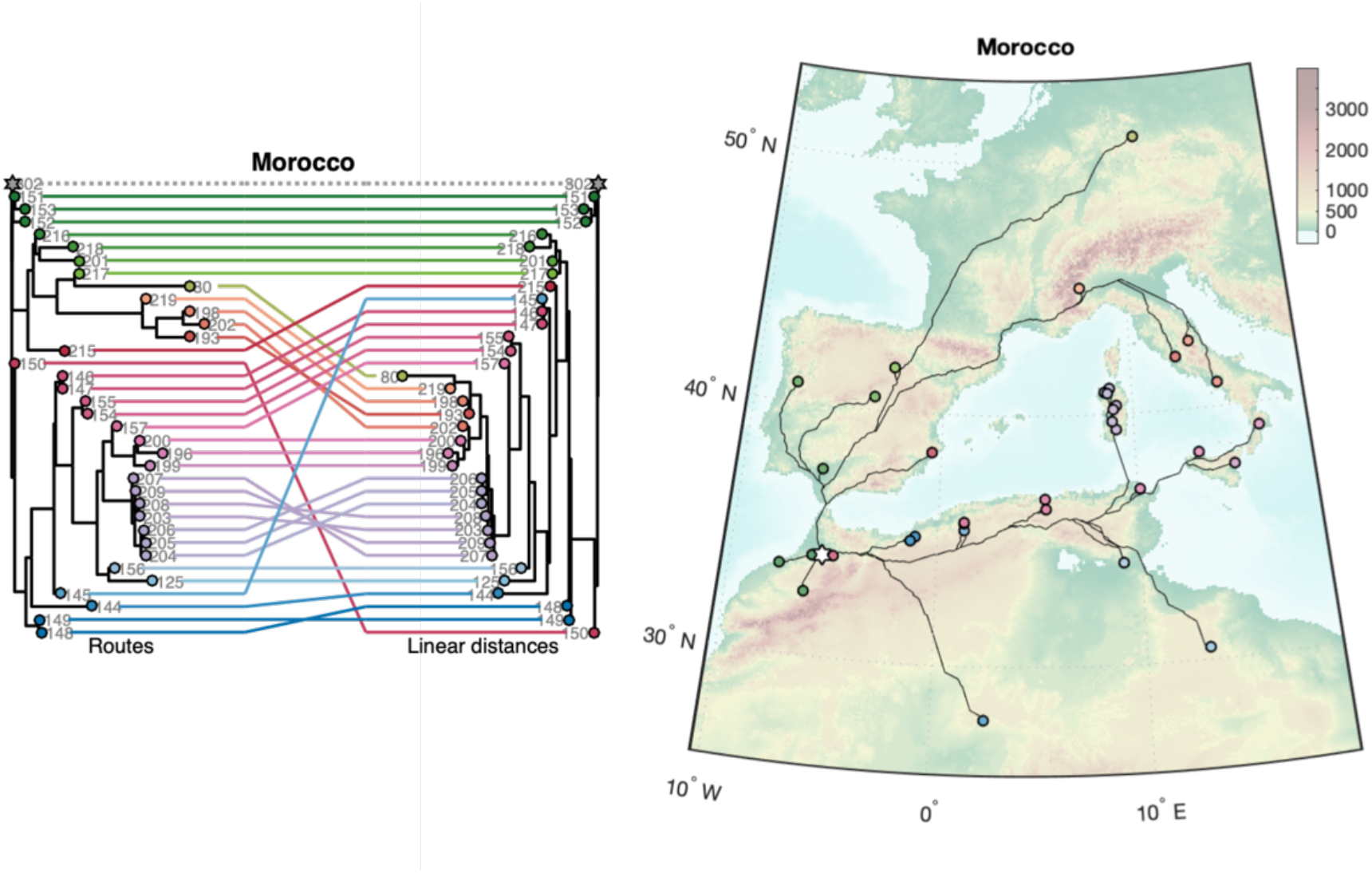
Tanglegram for correspondence between routes and linear distances within the Uzbekistan cluster. Routes of the Uzbekistan cluster on Map; star denotes the center of the cluster.

**Supplementary Figure 8.**
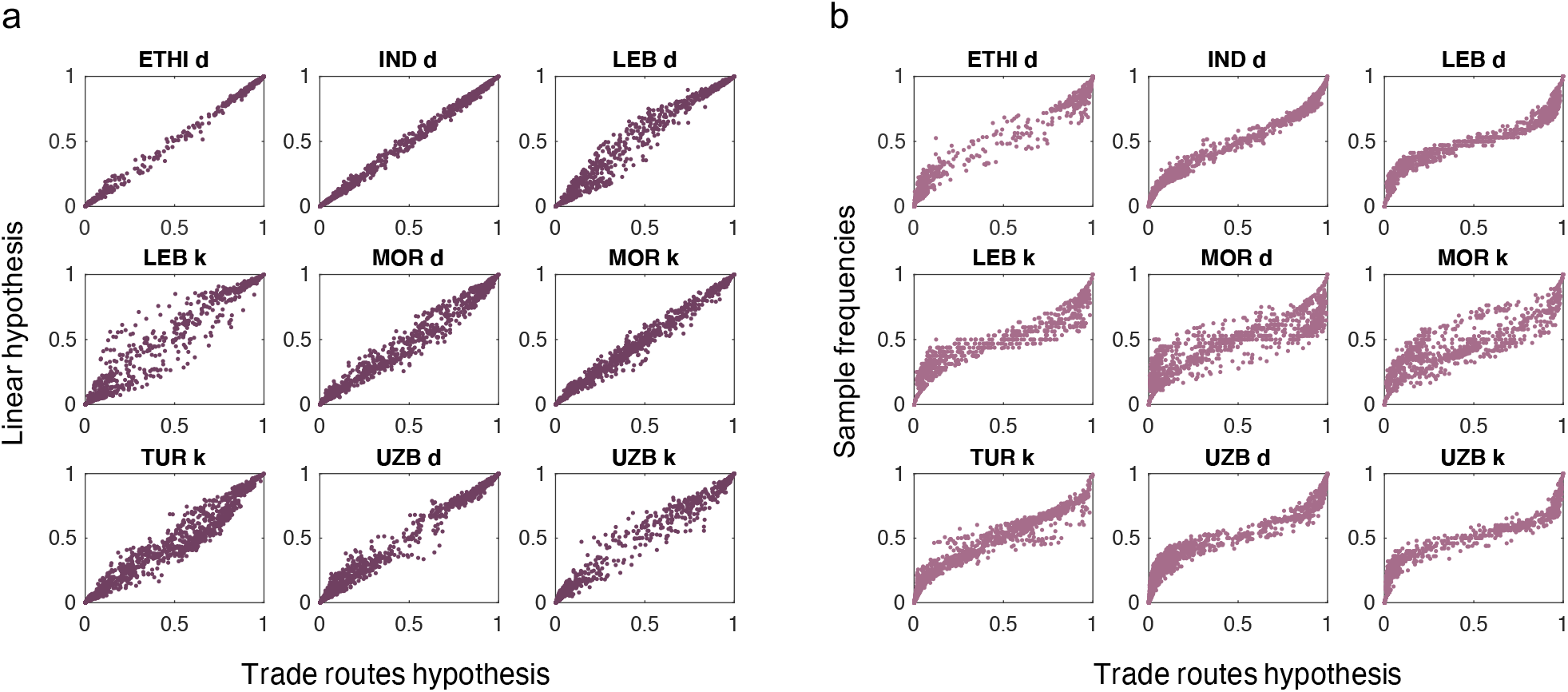
(a) Correspondence between allele frequencies estimated with **popdisp** under trade routes hypothesis and linear hypothesis. (a) Correspondence between allele frequencies estimated with **popdisp** under trade routes hypothesis and mean allele frequencies in populations.

**Supplementary Figure 9.**
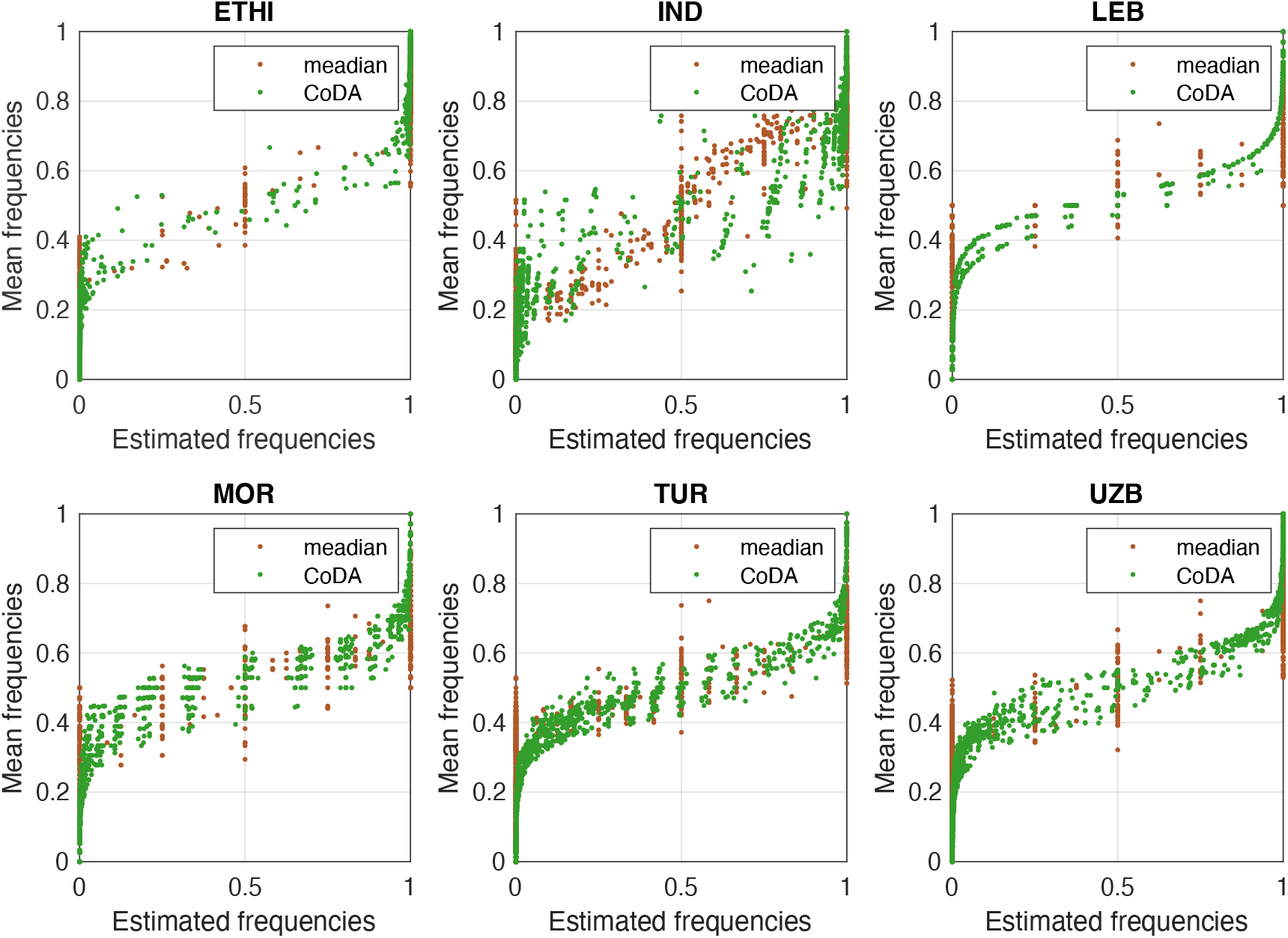
Correspondence between mean SNP frequencies in 6 desi populations and SNP frequencies estimated by two more robust methods. For each method, we took into account the regional distribution of samples: samples in each population belong to *n* geographical locations. For each SNP, we estimated the mean allele frequency in each location, 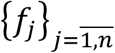, and then applied two methods. The first method (brown dots) reflects the median values across 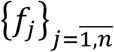. The second method (green dots) corresponds to the calculation of the center composition as in the compositional data analysis (CoDA). Together with mean allele frequencies in locations, this method considers frequencies of the second allele of the SNP, 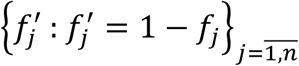. Then, it computes geometric mean on frequencies of each allele: 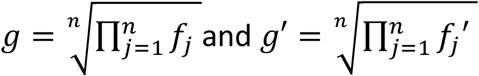. At last, it applies so-called closure function to obtained geometric means: 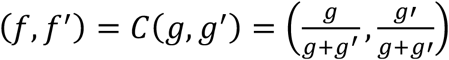 (Pawlowsky-Glahn and Buccianti 2011). Obtained *f* values for each SNP are “averaged” allele frequencies in a population in line with CoDA. Analysis of brown dots shows long vertical ranges at 0 and 1, indicating the prevalence of locations with homozygous SNPs, which is not caught by calculations of means. The CoDA-based method not only highlights the prevalence of SNP homozygosity but also softly accounts for minor heterozygosity. As our popdisp method, both methods (more robust than mean values) demonstrate S-like shape dependency between the mean and estimated SNP frequencies.

**Supplementary Figure 10.**
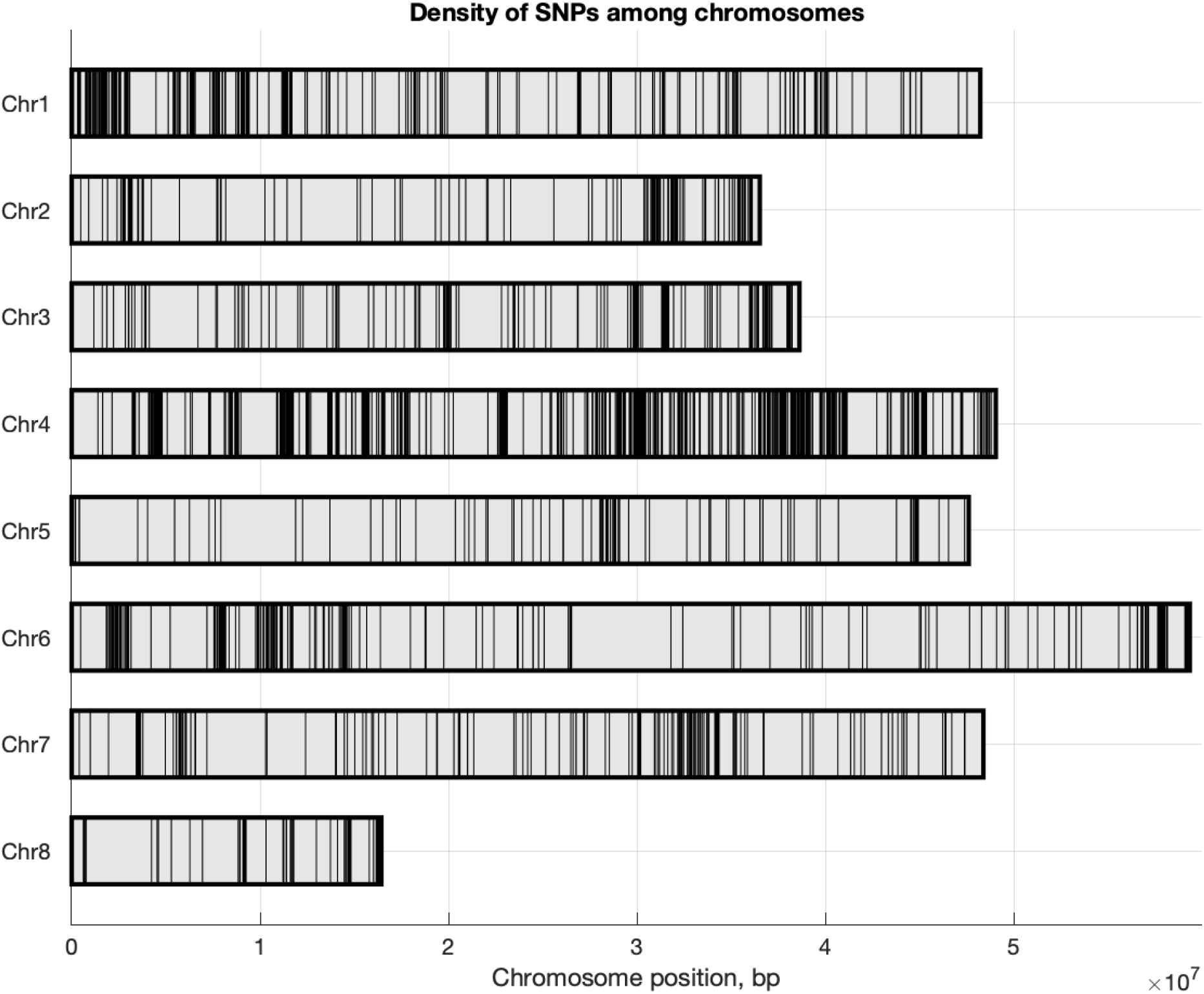
Density of SNPs along the chromosomes. Each vertical line corresponds to the position of one SNP.

